# Brachiopod genome unveils the evolution of the BMP–Chordin network in bilaterian body patterning

**DOI:** 10.1101/2024.05.28.596352

**Authors:** Thomas D. Lewin, Keisuke Shimizu, Isabel Jiah-Yih Liao, Mu-En Chen, Kazuyoshi Endo, Noriyuki Satoh, Peter W. H. Holland, Yue Him Wong, Yi-Jyun Luo

**Author notes:** Corresponding authors: Yue Him Wong, Yi-Jyun Luo.

## Abstract

Bone morphogenetic protein (BMP) signalling is crucial in regulating dorsal–ventral patterning and cell fate determination during early development in bilaterians. Interactions between BMP ligands and their main antagonist, Chordin, establish BMP gradients, subdivide embryos into distinct territories and organise body plans. However, the molecular control and evolutionary origins of dorsal–ventral patterning within spiralians, one of the three major bilaterian groups, have been obscured by their unique embryonic development. Here we present the chromosome-level genome of a spiralian with deuterostome-like development, the brachiopod *Lingula anatina*, and apply functional transcriptomics to study dorsal–ventral patterning under the control of BMP signalling. We uncover the presence of a dorsal–ventral BMP signalling gradient in the *L. anatina* gastrula with *bmp2/4* and *chordin* expressed at its dorsal and ventral sides, respectively. Using small-molecule drugs, exogenous recombinant BMP proteins and RNA sequencing, we show that a high level of BMP pathway activation inhibits the expression of neural genes during gastrula and larval stages. We also show that BMP signalling splits the developing larval shell field into two valves. The discovery of a BMP-mediated dorsal–ventral patterning system in a spiralian, similar to those observed in deuterostomes and non-spiralian protostomes, suggests deep conservation of this mechanism across all three major bilaterian clades. This is further supported by striking similarities in the gene sets regulated by BMP signalling in brachiopods and the vertebrate model *Xenopus*. We argue that the spiralian ancestor retained the basal bilaterian mechanism of dorsal–ventral patterning, although downstream components of the BMP–Chordin network have undergone dynamic evolutionary changes.

## Introduction

The integration of bone morphogenetic protein (BMP) signalling into the patterning of the dorsal–ventral axis is a cornerstone of bilaterian embryonic development. The common ancestor of bilaterians likely established this axis through an interplay between Bmp2/4 signalling molecules and their antagonists, such as Chordin^1–3^. In chordates, this interaction creates a BMP gradient, dividing the ectoderm into two distinct domains, a dorsal neural domain (low BMP) and a ventral epidermal domain (high BMP), in a process known as neural induction^4,5^. In the classical vertebrate model *Xenopus*, this gradient is regulated by the Spemann-Mangold organiser, a signalling centre that produces a cocktail of BMP antagonists, including Chordin^4,6,7^, Noggin^8,9^ and Admp^10^. In the fruit fly *Drosophila*, the BMP gradient is completely inverted, with dorsal Dpp/BMP and ventral Sog/Chordin, but neural domain formation is still induced at the low BMP end of the gradient^11–14^. This reversed pattern is considered to be the bilaterian ancestral state and is shared by protostomes and basal deuterostomes^15–17^. Aside from the dorsal–ventral axis inversion in chordates^16,18^, the ancestral BMP-mediated patterning mechanism remains highly conserved across diverse members of two of the three main groups of bilaterians: deuterostomes and ecdysozoans^19,20^.

Within Spiralia, a diverse group of protostomes including brachiopods, molluscs, annelids, and platyhelminths that constitutes the third major bilaterian clade, there appears to be no consistent role for BMP signalling in dorsal–ventral axis patterning^21^. As a sister group to Ecdysozoa, Spiralia displays diverse body patterning systems, including alternative BMP paralogues^22^, region-specific BMP signalling^23–25^, and even complete absence of BMP signalling^26^. For instance, perturbations to BMP signalling in the annelid *Capitella teleta* did not affect dorsal–ventral axis formation or central nervous system development but resulted in abnormal left–right asymmetries^24,27^. In the mollusc *Crepidula fornicata*, BMP signal perturbations affected neural and epidermal cell fates in the head region but not the trunk^23^. In another annelid, *Chaetopterus pergamentaceus,* Activin/Nodal signalling but not BMP signalling is the key specifier of the dorsal–ventral axis^26,28^. Indeed, the key BMP antagonist Chordin has been lost many times independently within annelids^29^ and other spiralians like platyhelminths^30^. In some species, such as the molluscs *Lottia goshimai* and *Ilyanassa obsoleta*, BMP signals involved in dorsal–ventral patterning are present but have a positive effect on neurogenesis^31,32^, opposite to that in ecdysozoans and deuterostomes. In light of these experiments, the ancestral function of the BMP signalling pathway in spiralian development remains unclear^27^. This highlights a major gap in understanding the evolution of the BMP–Chordin axis, particularly as spiralian embryos’ unique spiral cleavage^33,34^ complicates direct comparisons with other bilaterians at gastrulation, the key stage for dorsal– ventral patterning.

Brachiopods, especially those from lesser-studied benthic macrofauna in intertidal zones, have the potential to become invaluable models in studying BMP–Chordin axis evolution. Unlike typical spiralians, lingulid brachiopods exhibit radial embryonic cleavage^35^, possibly reflecting a reversion to the ancestral state, allowing direct gastrula stage comparisons with other deuterostome embryos. Previous studies have indicated that brachiopods such as *Novocrania anomala* and *Terebratalia transversa* may use a BMP– Chordin network for dorsal–ventral patterning that is similar to that of ecdysozoans and deuterostomes^36^. As a result, this group of brachiopods makes ideal subjects for comprehensive studies of the BMP–Chordin axis and for dissecting evolutionary scenarios between spiralians and other bilaterians.

Here we present the first chromosome-level genome of a brachiopod, *Lingula anatina*. We then incorporate this assembly into an updated phylogeny of lophotrochozoans and use comparative genomics to characterise BMP pathway component evolution in bilaterians. We further investigate dorsal–ventral patterning during *L. anatina* embryonic development. We find that BMP signalling is initially activated at the animal pole and subsequently at the dorsal side of the gastrula, suggesting deep conservation of the BMP gradient in spiralians despite their unique development. Using small-molecule drugs and recombinant proteins together with functional transcriptomics, we demonstrate that BMP signalling inhibits neuronal gene expression at both gastrula and larval stages. Our findings highlight the conserved aspects of the BMP–Chordin axis in brachiopods, reflecting similarities in body patterning with both protostomes and deuterostomes. This suggests that the spiralian ancestor employed a patterning mechanism similar to those found in these groups.

## Results

### Chromosome-level assembly of the brachiopod genome

To explore the functions of BMP signalling in the brachiopod *L. anatina*, we first sequenced its genome to chromosome level using PacBio HiFi (405-fold coverage) and Hi-C (171 million paired reads) technologies (Extended Data Fig. 1a–c and Supplementary Tables 1 and 2). The resulting assembly is 329.3 Mb in length with a 36.5% GC content and consists of 31.7% repetitive elements (Supplementary Fig. 1 and Supplementary Tables 3 and 4). There have been recent expansions of repeats, particularly long terminal repeat elements, in the genome of *L. anatina* and other Lophophorata members (Extended Data Fig. 2a,b). The assembly has a scaffold N50 of 30.1 Mb, a substantial improvement over the previous scaffold-level genome with a scaffold N50 of 0.5 Mb^37^, and is comparable in quality to other high-quality spiralian genomes^29,38–40^ (Fig. 1a and Supplementary Table 5).

**Fig. 1.**
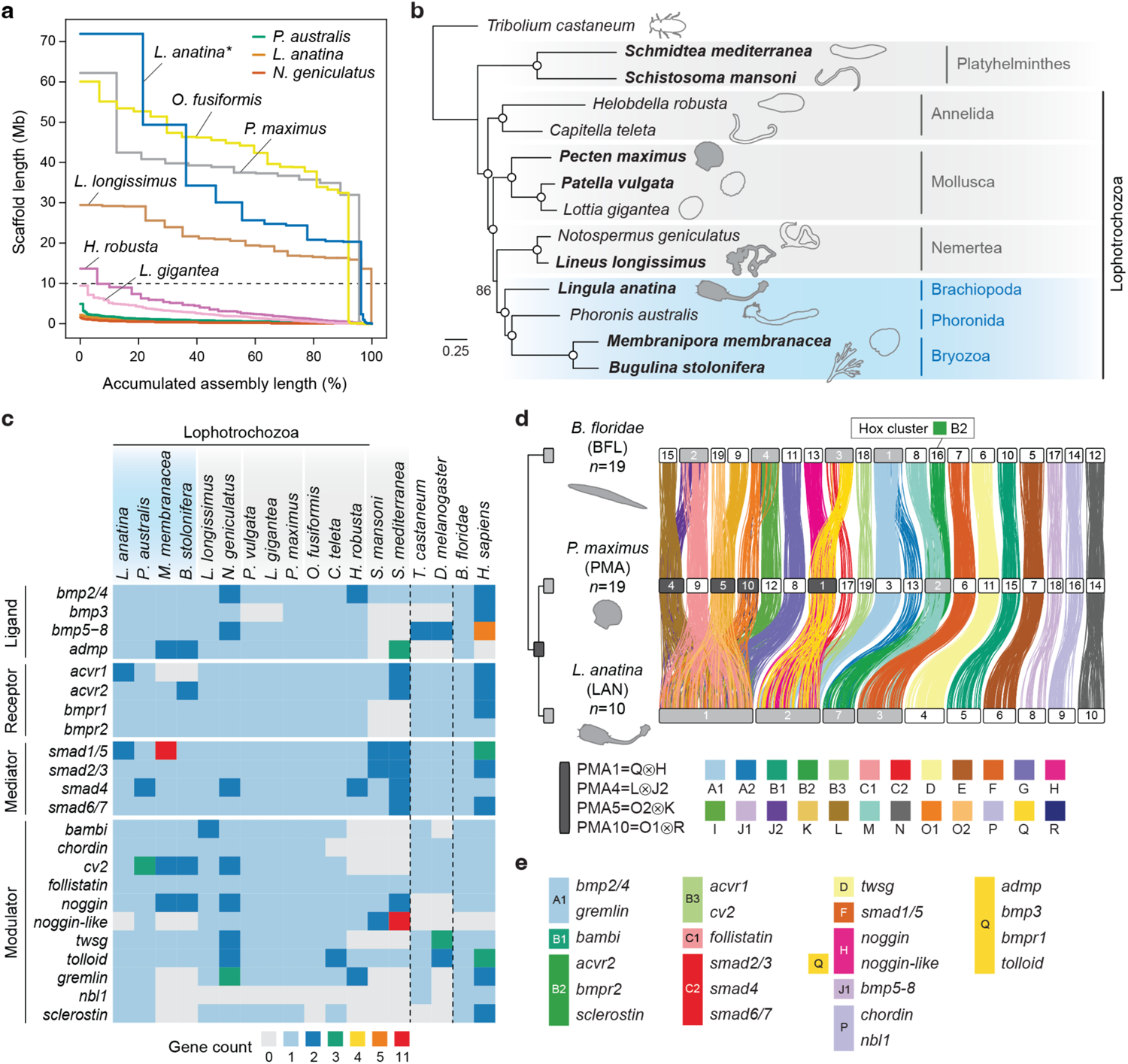
Brachiopod genome supports the Lophophorata hypothesis and reveals BMP signalling pathway evolution. **a**, Assembly quality comparison across selected spiralian genomes. The *L. anatina* assembly presented in this study (denoted with an asterisk) comprises ten chromosome-scale scaffolds. Horizontal dashed line at 10 Mb distinguishes scaffold-level genomes from chromosome-level genomes. **b**, Maximum likelihood phylogenetic inference of relationships between spiralians using the LG+F+R5 substitution model with 1,000 bootstrap replications. Phylogenetic tree was inferred from 109 single-copy orthologous genes with high topological robustness (average bootstrap support > 85%). Open circles denote 100% bootstrap support. Species in bold have chromosome-level assemblies. Species with shaded silhouettes are included in further synteny analysis. The traditional Lophophorata clade, highlighted in blue, comprises Brachiopoda, Phoronida and Bryozoa. **c**, Heatmap showing BMP signalling pathway gene repertoires in bilaterians. Vertical dashed lines separate the three main bilaterian clades: spiralians, ecdysozoans and deuterostomes. Lophophorata is shaded in blue. **d**, Chromosome-scale gene linkage observed between amphioxus, scallops and brachiopods. Horizontal bars denote chromosomes. Dark grey chromosomes indicate ancestral spiralian fusion events, while light grey chromosomes represent lineage-specific fusion events. Vertical lines between chromosomes link the genomic positions of orthologous genes and are colour-coded based on bilaterian ancestral linkage groups (ALGs)^41^. The *L. anatina* genome has undergone extensive rearrangements, with lineage-specific fusion events resulting from chromosomes 1, 2, 3, and 7. Notably, the expansive *L. anatina* chromosome 1 equates to six distinct chromosomes in *P. maximus*. In the selected bilaterians, the Hox cluster’s location aligns with its affiliation to ALG B2. **e**, Conserved associations of BMP signalling pathway genes and ALGs in amphioxus, scallops and brachiopods.

The majority of the assembly consists of 10 chromosome-scale scaffolds accounting for 97.8% of the total length, consistent with previous karyotype observations (*n* = 10)^42^. Gene prediction and annotation using newly generated RNA sequencing data resulted in highly complete gene models with a BUSCO score of 98.1% (Metazoa *odb10*) (Extended Data Fig. 1d and Supplementary Tables 6–10). These gene models comprise 29,458 transcripts corresponding to 24,330 unique protein-coding genes, with extensive redundancy reduction achieved compared to the previous assembly (Supplementary Fig. 1 and Supplementary Table 3). Gene family analysis reveals a gene content that is more stable than those of other Lophotrochozoans (Supplementary Fig. 2 and Supplementary Table 11). Notably, we found that longer chromosomes have a higher density of protein-coding genes and a lower density of repeats (Extended Data Fig. 3).

### Spiralian phylogeny and the Lophophorata hypothesis

Reliable phylogenetic relationships are essential for understanding the direction of evolutionary change^43^, yet the phylogenetic relationships between spiralians remain highly contentious^44–46^. To explore the evolution of the BMP–Chordin pathway, we built an updated phylogenetic tree focussed on the Lophotrochozoa, a major clade within Spiralia, by integrating newly sequenced chromosome-level genomes of brachiopods (this study), bryozoans^47,48^, nemerteans^40^, and molluscs^39,49^ (Supplementary Table 12). We then selected only orthologues with the strongest signal for phylogenetic reconstruction (see Methods). Using this method, we identified a monophyletic Lophophorata, in which Bryozoa and Phoronida form a clade that is sister to Brachiopoda (Fig. 1b). Historically, the shared morphological feature of the lophophore—a ciliated tentacle-like feeding structure—has supported this clade, but several molecular studies have questioned the inclusion of bryozoans in this taxonomic grouping^50–52^. Intriguingly, we found that species trees constructed from orthologues with weaker phylogenetic signals support a polyphyletic Lophophorata (Extended Data Fig. 4). Our analysis suggests that the Polyzoa topology, which groups Bryozoa with Entoprocta and Cycliophora^51^, may emerge when unreliable data with a high signal-noise ratio is used.

### BMP pathway components in bilaterian genomes

The absence of high-quality genomes has hindered accurate assessment of gene duplications and losses in numerous spiralian groups, especially in understudied lineages such as brachiopods, phoronids, bryozoans, and nemerteans. To reconstruct the evolution of the BMP signalling pathway in bilaterians, we utilised several lophotrochozoan and outgroup species with chromosome-level assemblies. We then annotated BMP pathway components in these genomes based on reported pathway genes in model systems such as *Xenopus*^5^, *Drosophila*^53^ and sea urchins^54^ and sought to characterise the evolution of BMP components through gene duplication and loss analysis (Fig. 1c, Supplementary Fig. 3 and Supplementary Tables 13 and 14). We found that the main ligands and modulators associated with dorsal– ventral patterning, notably *bmp2/4* and *chordin*, are highly conserved in the Lophotrochozoa, with *bmp2/4* present in all sampled species and *chordin* only lost within annelids^29^ (Fig. 1c).

In contrast, other BMP components show remarkable variation considering their developmental importance. This is epitomised by bryozoans, which possess numerous lineage-specific duplications and losses to many key components at a scale comparable to known divergent groups like platyhelminths^55^. Two copies each of *admp*, *cv2* and *noggin* are present and we failed to find *acvr1*, *noggin-like* and any of the three *DAN* family genes. We also found 11 copies of *smad1/5* in the bryozoan *Membranipora membranacea*. Dynamic duplications and losses are not limited to bryozoans: the brachiopod *L. anatina* has a *smad1/5* duplicate, a partial *acvr1* duplicate and *noggin-like* is absent; the phoronid *Phoronis australis* has two copies of *smad4* and three of *cv2*; the nemertean *Lineus longissimu*s has two copies of *bambi*; two mollusc assemblies lack *bmp3*; two annelid assemblies lack *sclerostin*; and *nbl1* is missing from all selected species except *L. anatina* and *P. australis*. Numerous gene duplicates are identified in the nemertean *Notospermus geniculatus*, but based on BUSCO duplication scores (Extended Data Fig. 1) and the lack of duplicates in the nemertean *L. longissimus*, these are likely attributable to false haplotypic duplications^56,57^. This finding highlights the need for high-quality chromosome-level genomes for gene content analyses. Overall, BMP pathway repertoires exhibit notable evolutionary variation within spiralians, indicating that BMP gradients are intricately modulated and may have evolved to adapt diverse developmental modes and environmental contexts across various bilaterian lineages.

### Conserved syntenic associations of BMP pathway genes

We next questioned whether the developmental importance of BMP genes as an ancestral mechanism for body patterning in bilaterians has affected the evolution of their genomic position. To explore this hypothesis, we analysed genome-scale linkage conservation, focusing on the macrosynteny of BMP genes in five selected bilaterians: the brachiopod *L. anatina*, the mollusc *Pecten maximus*^39^, the nemertean *L. longissimus*^40^, the annelid *Owenia fusiformis*^29^ and the chordate *Branchiostoma floridae*^58^. We first identified the correspondence between chromosomes and bilaterian ancestral linkage groups (ALGs)^41^, then characterised the events of chromosome fusion and fission (Fig. 1d and Supplementary Tables 15 and 16). Strikingly, our analysis reveals massive lineage-specific fusion-with-mixing in *L. anatina*, with chromosomes 1, 2, 3 and 7 all the products of fusion events (Fig. 1d). The giant 71.9 Mb *L. anatina* chromosome 1 consists of nine bilaterian ALGs (C1, G, I, J2, K, L, O1, O2 and R) and corresponds to six separate *P. maximus* chromosomes (Extended Data Fig. 5). Further comparison with the annelid *O. fusiformis* and nemertean *L. longissimus* suggests independent fusion events in brachiopods and annelids (Extended Data Fig. 6). We next identified the genomic locations of BMP pathway components to assess their associations with an ALG. Our results show remarkable conservation: in each of the five selected bilaterian genomes, all 23 annotated BMP components are invariably linked to the same ALG (Fig. 1e and Supplementary Table 17). This exceeds the usual ALG conservation rate found in randomly selected genes (chi-square test, *p* = 0.003) (Supplementary Table 18), suggesting a higher tendency for BMP pathway genes to maintain their specific ALG associations. Evolutionary conservation of the genomic locations of BMP pathway components may reflect the preservation of regulatory programs across large evolutionary spans.

To determine if the conservation of ALG associations is a characteristic unique to BMP pathway components, we extended our analysis to other genes with conserved, ancient roles in development. Remarkably, all 12 Wnt ligands found in both spiralians and chordates demonstrated 100% association with a single ALG across the five studied species, a rate significantly higher than the average ALG conservation (chi-square test, *p* = 0.038) (Supplementary Table 19). In addition, the Hox gene cluster is consistently associated with bilaterian ALG B2 in all analysed species. This persistent conservation of developmental genes’ genomic locations, from brachiopods to chordates, despite widespread chromosomal rearrangements, implies evolutionary constraints on the mobility of these genes among ALGs. We propose that there is a stronger selective pressure at crucial developmental loci to maintain associations with regulatory elements in nearby genomic regions, thereby preventing their translocation and resulting in highly conserved macrosynteny.

### Dynamic expression of BMP pathway genes and BMP gradients

The co-option of BMP–Chordin signalling pathway into dorsal–ventral axis patterning is a cornerstone of the bilaterian body plan (Fig. 2a). This pathway operates through Smad1/5 phosphorylation, leading to nuclear translocation and regulation of downstream BMP signalling genes (Fig. 2b). To explore BMP–Chordin pathway gene expression in *L. anatina*, we quantified the expression of BMP pathway genes in RNA-sequencing data^37^ (Extended Data Fig. 7 and Supplementary Table 20). We found that *smad1/5* and *bmp5–8* are expressed maternally, with their expression persisting into the larval stages (Fig. 2c). Key BMP genes *bmp2/4* and *chordin* show early upregulation in the blastula, while modulators *bambi* and *cv2* increase during the gastrula stage. BMP receptors are ubiquitously expressed, indicating a non-crucial role in the timing of BMP pathway activation (Extended Data Fig. 8). Interestingly, modulators like *follistatin*, *gremlin*, *noggin* and *nbl1* show no expression in these stages, suggesting that they are not involved in dorsal–ventral patterning.

**Fig. 2.**
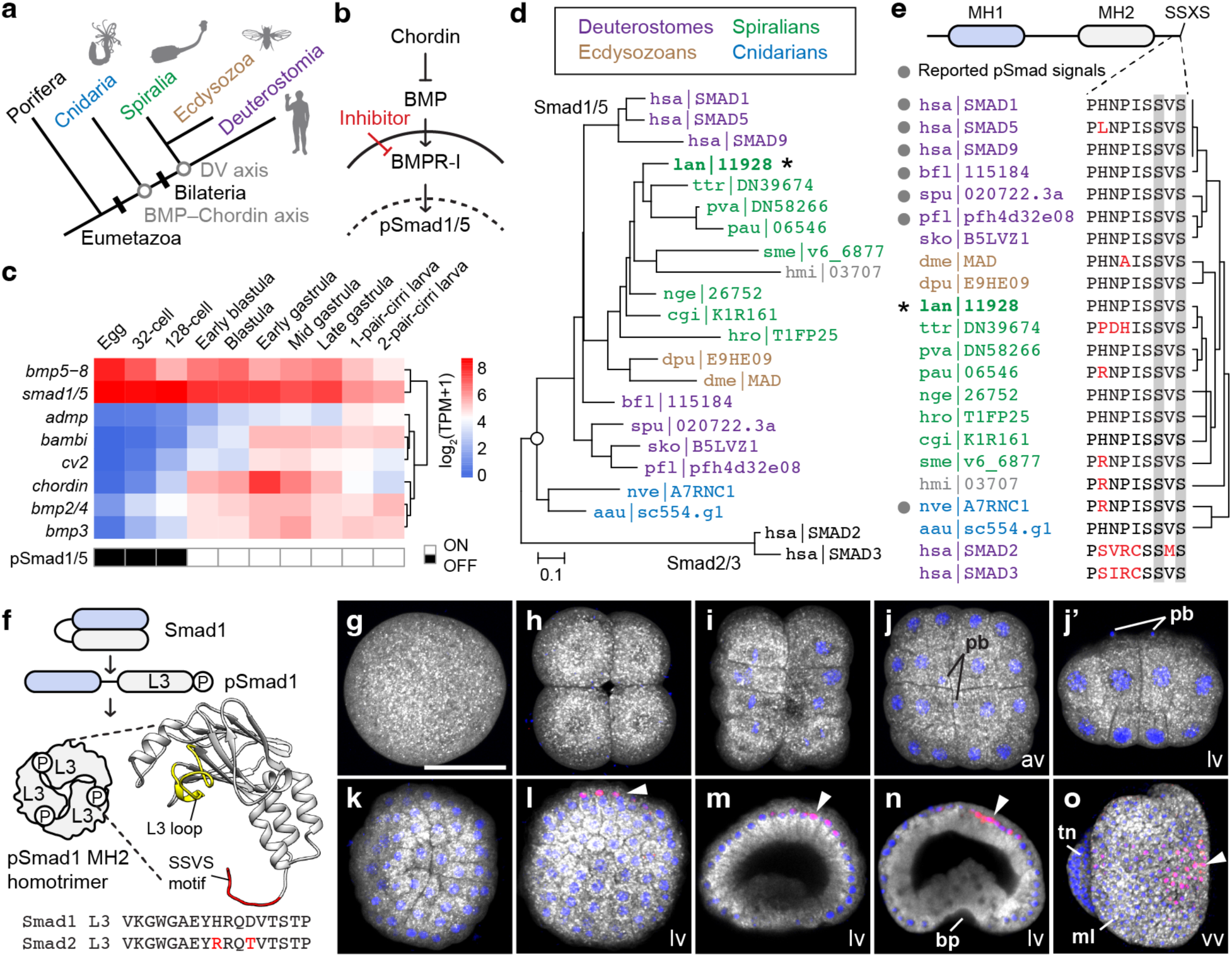
Brachiopod embryos express all essential BMP pathway components and exhibit a BMP gradient. **a**, Schematic illustration showing the evolution of the BMP– Chordin axis in animals. The BMP-Chordin axis predates the origin of bilaterians and, therefore, the dorsal–ventral (DV) axis. **b**, Canonical BMP signalling pathway regulated by Chordin and mediated through BMP type I receptor (BMPR-I) and phosphorylated Smad1/5 (pSmad1/5). Solid line represents the plasma membrane, while dashed line denotes the nuclear envelope. **c**, Expression profiles of *L. anatina* main BMP signalling ligands, mediators, and modulators. Appearance of nuclear pSmad1/5 signals is shown in blank rectangles. TPM, transcripts per million. **d**, Phylogeny of receptor-regulated Smads (R-Smads) inferred with 22 genes (427 amino acid positions) using the maximum likelihood method with the LG model and 1,000 bootstrap replications. Numbers at the nodes indicate bootstrap support values. Proteins are identified by their UniProt, gene model, or transcriptome IDs. *L. anatina* Smad1/5 is indicated with an asterisk. **e**, Upper: Domain composition of an R-Smad protein (400–500 amino acids). MH, mother against dpp homology. Lower: Alignment of C-terminus of R-Smad proteins. Phosphorylated sites corresponding to Ser463/465 in human Smad1 are shaded in grey. Different amino acids compared to Smad1 are labelled in red. Grey circles indicate reported pSmad1/5 signals using the antibody against the phosphopeptide PHNPISSVS. Species tree on the right. **f**, Schematic illustration of R-Smad activation. MH1 domain for DNA binding is shown in blue, and MH2 domain for interacting with other Smads, such as Smad4, is shown in grey. Model of *L. anatina* Smad1/5 MH2 structure aligned to human Smad1 MH2 domain (Protein Data Bank entry 1KHU). Conserved L3 loop and SSVS motif are coloured yellow and red, respectively. The L3 loop contains 17 amino acids (VKGWGAEYHRQDVTSTP). Different amino acids compared to Smad1 are labelled in red. **g***–***o**, Immunostaining of pSmad1/5 (red) at early embryonic stages shows signals with asymmetrical nuclear localisation (arrowheads). Nuclei are labelled with Hoechst 33342 (blue). Cytosol is counterstained with CellMask Deep Red (grey). Embryonic stages: 1-cell (**g**); 4-cell (**h**); 16-cell (**i**); 32-cell (**j** and **j’**); 64-cell (**k**); blastula (**l**); early gastrula (**m**); late gastrula (**n**); early larva, ventral to the left (**o**). pb, polar body; bp, blastopore; ml, mantle lobe; tn, tentacle; av, animal view; lv, lateral view; vv, ventral view. Scale bar, 50 μm. Three-letter species code: hsa, human (*Homo sapiens*); bfl, amphioxus (*Branchiostoma floridae*); spu, sea urchin (*Strongylocentrotus purpuratus*); pfl, hemichordate (*Ptychodera flava*); sko, hemichordate (*Saccoglossus kowalevskii*); dme, fruit fly (*Drosophila melanogaster*); dpu, water flea (*Daphnia pulex*); lan, brachiopod (*Lingula anatina*); ttr, brachiopod (*Terebratalia transversa*); pva, phoronid (*Phoronis vancouverensis*); pau, phoronid (*Phoronis australis*); nge, nemertean (*Notospermus geniculatus*); hro, leech (*Helobdella robusta*); cgi, Pacific oyster (*Crassostrea gigas*); sme, planarian (*Schmidtea mediterranea*); hmi, acoel (*Hofstenia miamia*); nve, sea anemone (*Nematostella vectensis*); aau, jellyfish (*Aurelia aurita*).

The Smad1/5 protein, pivotal to BMP signalling across metazoans (Fig. 2d), possesses MH1 and MH2 domains and an SSXS motif phosphorylation site essential for DNA binding and protein trimerisation (Fig. 2e,f). We found that, although the *L. anatina* genome contains two copies of the *smad1/5* gene, only one of these copies is active during early development, while the other is most highly expressed in the gonad (Extended Data Fig. 8 and Supplementary Fig. 4). Using an antibody targeting the conserved PHNPISSVS phosphopeptide sequence, we examined the distribution of phosphorylated Smad1/5 (pSmad1/5) as an indicator of BMP signalling activity. We found that nuclearised pSmad1/5 is absent from the 1-cell to 64-cell stages (Fig. 2g–k) but appears in the blastula, asymmetrically localised at the animal pole (Fig. 2l). This occurrence aligns with the upregulation of *bmp2/4*, *bmp3* and *chordin* (Fig. 2c), suggesting their importance in early BMP signalling regulation. The presence of pSmad1/5 then persists on the dorsal side through the gastrula and early larval stages (Fig. 2m–o), demonstrating dorsal BMP activation. This pattern is consistent with those found in other protostomes and basal deuterostomes, underscoring a deeply conserved BMP gradient in spiralians.

### BMP gradients regulate dorsal–ventral patterning

To investigate the role of the BMP signalling pathway’s asymmetric activation, we applied BMP receptor inhibitors and recombinant BMP proteins for two durations: from early blastula to late gastrula and from early blastula to the one-pair-cirri larva (Fig. 3a and Supplementary Table 21). Control late gastrula embryos immunostained for pSmad1/5 showed expected dorsal localisation (Fig. 3b,c). However, embryos treated with the BMP receptor inhibitors LDN-193189 (LDN)^59^ and K02288 (K02)^60^, collectively referred to as BMP(-), showed a complete absence of pSmad1/5 staining (Fig. 3d–e). Conversely, treatment with recombinant mouse BMP4 (mBMP4) protein^61^, referred to as BMP(+), led to a loss of dorsal pSmad1/5 localisation and an increase in pSmad1/5-positive nuclei across the gastrula (Fig. 3f), validating the impact of our manipulations on BMP signalling. Notably, BMP(-) and BMP(+) treatment also altered gastrula morphology (Fig. 3c–e), including by delaying gastrulation, emphasising the significance of the BMP gradient for gastrula development.

**Fig. 3.**
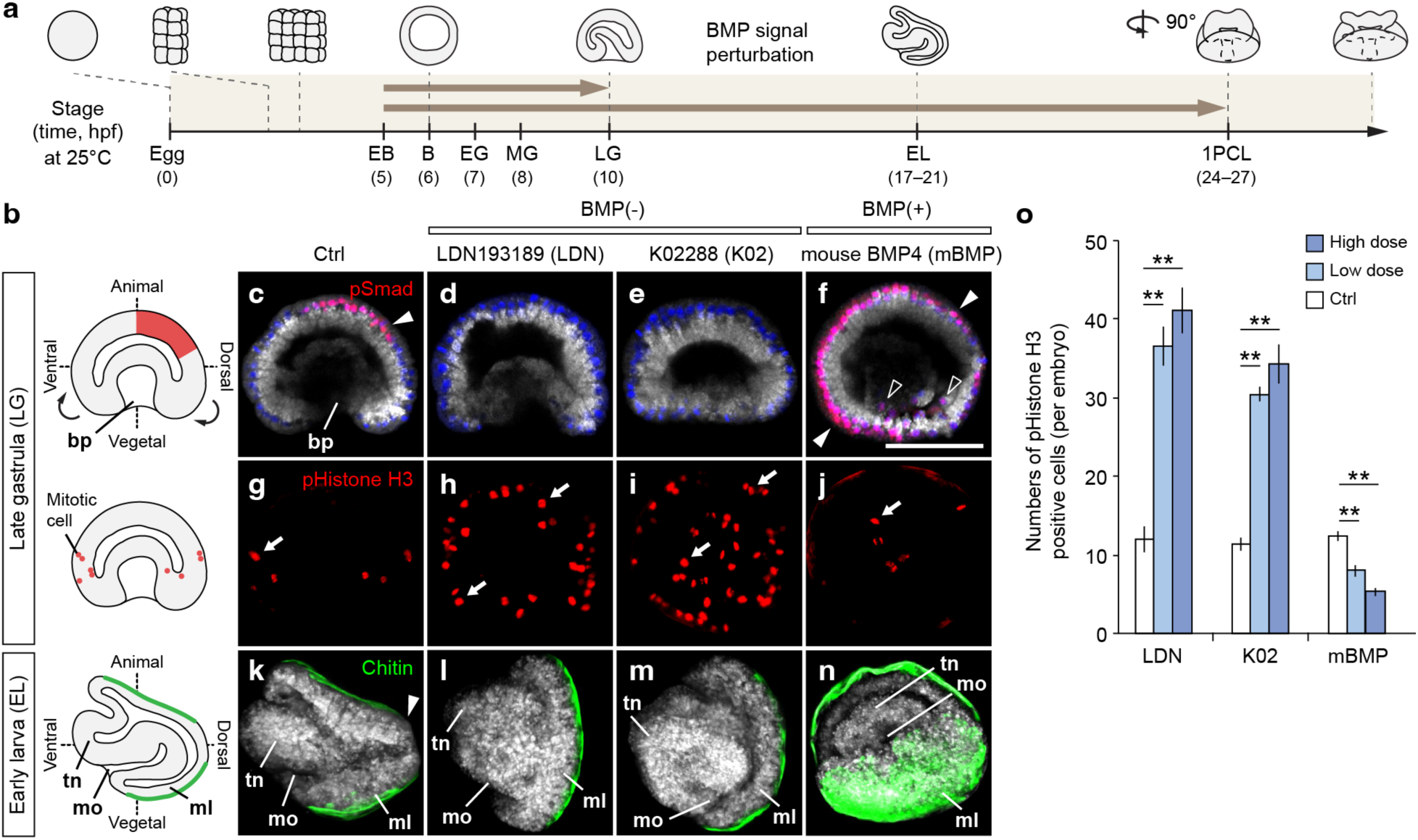
BMP gradients regulate brachiopod dorsal–ventral patterning and cell proliferation. **a**, Schematic illustration of *L. anatina* developmental timeline (hours post-fertilisation, hpf) and manipulation of BMP signalling. The embryos were treated with inhibitors (LDN193189 and K02288) or recombinant proteins (mouse BMP4) for two incubation windows (brown arrows). EB, early blastula; B, blastula; EG, early gastrula; MG, mid-gastrula; LG, late gastrula; EL, early larva; 1PCL, one-pair-cirri larva. 1PCL displays a view rotated 90 degrees from the plane relative to EL. **b**, Schematic diagram of *L. anatina* embryos with labelled body axes. bp, blastopore; tn, tentacle; mo, mouth; ml, mantle lobe. Curved arrows show the cell movement at the gastrula stage, giving rise to the early larva. A BMP gradient labelled with pSmad1/5 is shown as a red region. Mitotic cells labelled with phospho-Histone H3 (pHistone H3) are shown in red circles. Chitinous embryonic shells are shown in green. **c**–**f**, Immunostaining of pSmad1/5 antibody (red) in optical sectioned late gastrulae. Nuclei are labelled with Hoechst 33342 (blue). Cytosol is counterstained with CellMask Deep Red (grey). Nuclearised pSmad1/5 signals in red (overlap of pSmad1/5 and Hoechst 33342 signals) are indicated by arrowheads. Scale bar, 50 μm. **g**–**j**, Immunostaining of pHistone H3 antibody (red). Mitotic cells are indicated by arrows. **k**–**n**, Staining of chitin with a chitin-binding probe (green). Chitin staining marks the embryonic shells. Note that the dorsal edge of the larva is absent of chitin staining in the control (**k**). **o**, Statistics of cell proliferation under BMP signalling perturbation (*n* = 5). High dose (LDN, 8 μM; K02, 500 nM; BMP 200 ng/mL); low dose (LDN, 4 μM; K02, 250 nM; BMP 100 ng/mL); error bars, standard errors of the mean; double asterisks, *t*-tests for comparing between control and treatments (*p* < 0.01).

We next explored BMP signalling’s influence on cell proliferation by counting gastrula cells marked with the mitosis-specific phospho-histone H3. We observed that BMP(-) treatments significantly increased proliferative cell numbers compared to the control, whereas BMP(+) embryos showed a decrease in mitotic cells (*p* < 0.01) (Fig. 3g–j,o). This indicates that one of the roles of BMP signalling is to inhibit cell proliferation in *L. anatina* gastrulae. During *L. anatina* development, the anterior–posterior axis forms dorsal and ventral mantle lobes through mantle lobe folding (Fig. 3b)^35^. We assessed the necessity of BMP signalling to this process by manipulating the BMP pathway and observing embryonic shell formation with a chitin-binding probe. In control embryos, dorsal–ventral folding is marked by chitin staining across both mantle lobes, excluding the dorsal edge (Fig. 3k). In contrast, BMP(-) embryos showed disrupted dorsal–ventral folding, indicated by continuous chitin staining, including at the dorsal edge (Fig. 3l–m). BMP(+) embryos exhibited ’over-folding’, with the chitinous shell extending ventrally, resulting in the tentacle and mouth being encased inside the shell (Fig. 3n). These findings indicate that dorsal BMP signalling is essential for dorsal–ventral folding and splitting the developing larval shell into two distinct valves. In summary, our results show that BMP signalling, highest at the dorsal side in *L. anatina* as in other protostomes and basal deuterostomes, is crucial for regulating cell division and maintaining embryonic structural integrity.

### BMP signals control neuronal and cell cycle gene expression

We further explored the role of BMP signalling in *L. anatina* development through functional transcriptomics, using RNA sequencing on control, BMP(-) and BMP(+) embryos (Supplementary Tables 22 - 24). Analysis of differentially expressed genes at the gastrula stage revealed differential transcriptomic changes: 881 genes showed strongly significant expression changes (fold-change > 4, *p* < 0.001) under BMP manipulations, as identified by hierarchical clustering (Fig. 4a–d). Gene ontology (GO) analysis (Supplementary Tables 25 - 28) highlighted ’nervous system development’ as a highly enriched term among downregulated genes (Fig. 4e,f). Focusing on a robust dataset of genes differentially expressed at both gastrula and early larval stages (49 BMP-upregulated and 45 BMP-downregulated genes) (Fig. 4g,h), we found the BMP-downregulated targets to be dominated by neuronal genes, including *alk*, *elav*, *foxb1*, *lhx1/5*, *netrin*, *nkx2.4*, *otx2*, *pax6*, *six3/6*, *sim1/2*, *soxB1* and *soxB2*^62–69^ (Fig. 4h). This finding contrasts with a recent study that highlights BMP’s positive influence on neurulation in spiralians^31,32^, but is consistent with the observed inhibitory effect of BMP in the neuroectoderm of non-spiralian protostomes^70^ and deuterostomes^16,20^. Our results suggest that inhibition of neural development by elevated BMP levels is a conserved mechanism among spiralians, despite the evolutionary diversification of embryological development modes in molluscs^31^ and annelids^24^. In addition, at the gastrula stage, we found ‘DNA replication’, ‘chromosome organisation’ and ‘cell cycle’ as the most statistically significant GO terms for downregulated genes. This is consistent with the above result that the presence of mitotic cells was reduced in BMP(+) embryos and increased in BMP(-) embryos.

**Fig. 4.**
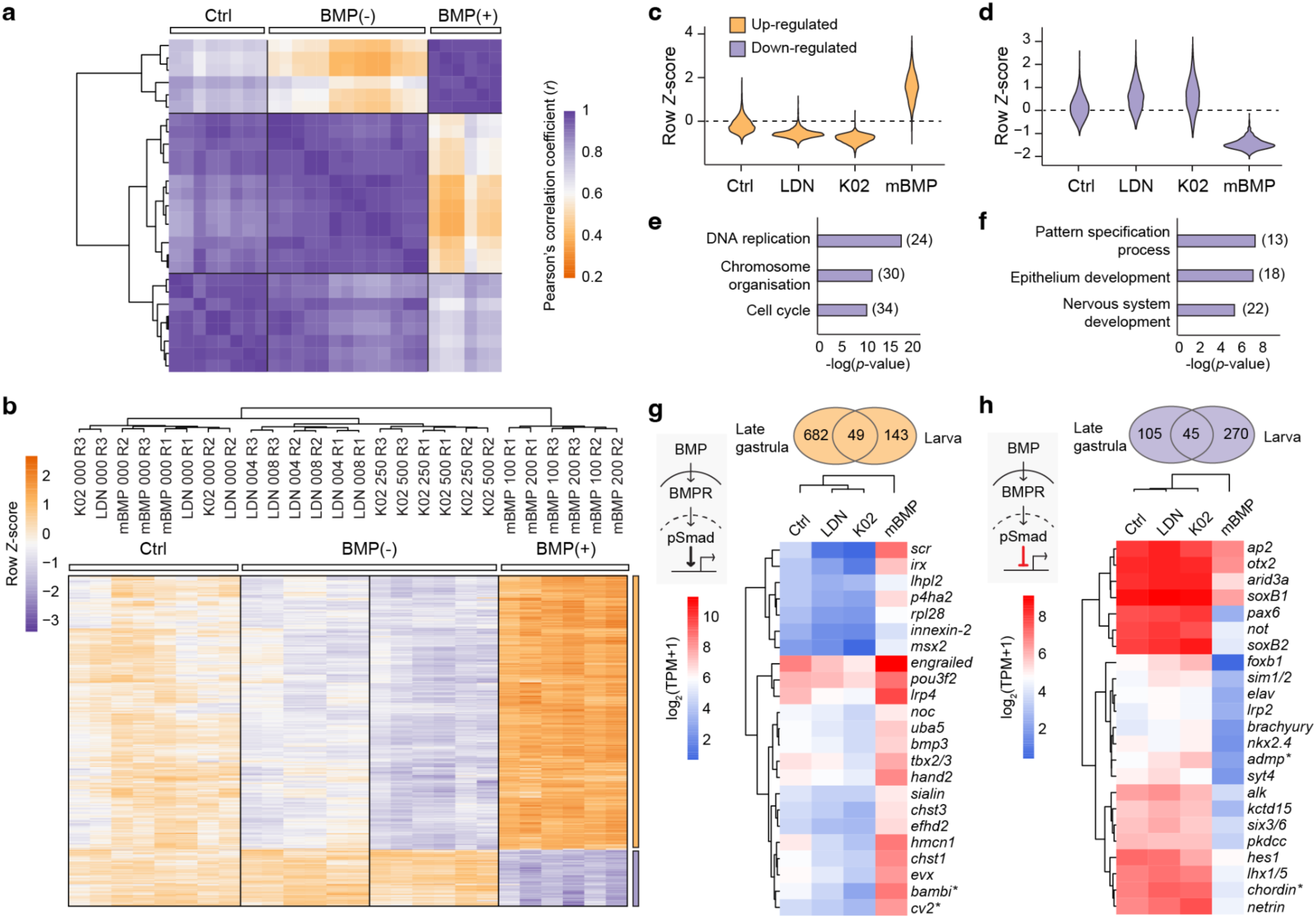
BMP signals suppress the expression of genes involved in neurogenesis. **a**, Pearson’s correlation analysis based on 881 differentially expressed genes (fold-change > 4, *p* < 0.001) under the manipulation of BMP signals. Each row and column represents one sample. Pearson’s correlation coefficient (*r*) measures the similarity of samples’ transcriptomes: purple being the highest similarity and orange the lowest. **b**, Hierarchical clustering heat map of 881 differentially expressed genes at the late gastrula stage under the manipulation of BMP signals. The *Z*-score scale represents log-transformed and mean-subtracted transcripts per million (TPM). BMP up-regulated (731, orange) and down-regulated (150, purple) gene groups are labelled on the right. Treatments and units: LDN193189 (LDN), μM; K02288 (K02), nM; mouse BMP4 (BMP), ng/mL. R1–R3, biological replications. **c** and **d**, Violin plots showing the expression distribution of up-regulated (orange) (**c**) and down-regulated (purple) (**d**) gene groups under treatments. **e** and **f**, Gene ontology enriched biological process terms at the late gastrula stage (**e**) and early larval stage (**f**) that are down-regulated (purple) under the manipulation of BMP signalling. Numbers of genes within the functional groups are shown in parentheses. **g** and **h**, Venn diagrams: Differentially expressed genes shared by late gastrula and early larval stages. Heatmaps: Expression profiles of selected BMP up-regulated (**g**) and down-regulated (**h**) genes at the late gastrula stage. Asterisks highlight *Xenopus* ventral centre and Spemann-Mangold organiser genes.

### Deep conservation of the dorsal–ventral patterning pathway within bilaterians

We next investigated whether the downstream effects of BMP signalling on gene expression that we observed in *L. anatina* show similarities to those of other bilaterians. We selected the vertebrate model *Xenopus* for this comparison, as the role of BMP signalling in dorsal–ventral patterning is well characterised^5^ and its phylogenetic position within the deuterostomes means that evolutionary inferences can be made that span the entire Bilateria. In *Xenopus*, a ventral signalling centre produces the BMP ligand Bmp4^71^ plus feedback inhibitors like Cv2^72,73^, Twisted-gastrulation^74–76^ and Bambi^77^, while the dorsal Spemann-Mangold organiser secretes a set of Bmp antagonists which includes Chordin^4,6,7^, Follistatin^78^, Noggin^8,9^ and Admp^10^. These antipodal signalling centres construct the dorsal–ventral BMP gradient that patterns the axis, with the neural inducers secreted from the Spemann-Mangold organiser causing the formation of neural tissue on the dorsal side. We found that several brachiopod homologues of key *Xenopus* dorsal–ventral patterning genes were up-or down-regulated at the RNA level in the above BMP signal manipulation experiments. Specifically, BMP signalling in *L. anatina* downregulates homologues of the *Xenopus* Spemann-Mangold organiser genes *chordin* and *admp*, while upregulating homologues of the *Xenopus* ventral centre genes *bambi* and *cv2*. (Fig. 4g,h).

**Fig. 5.**
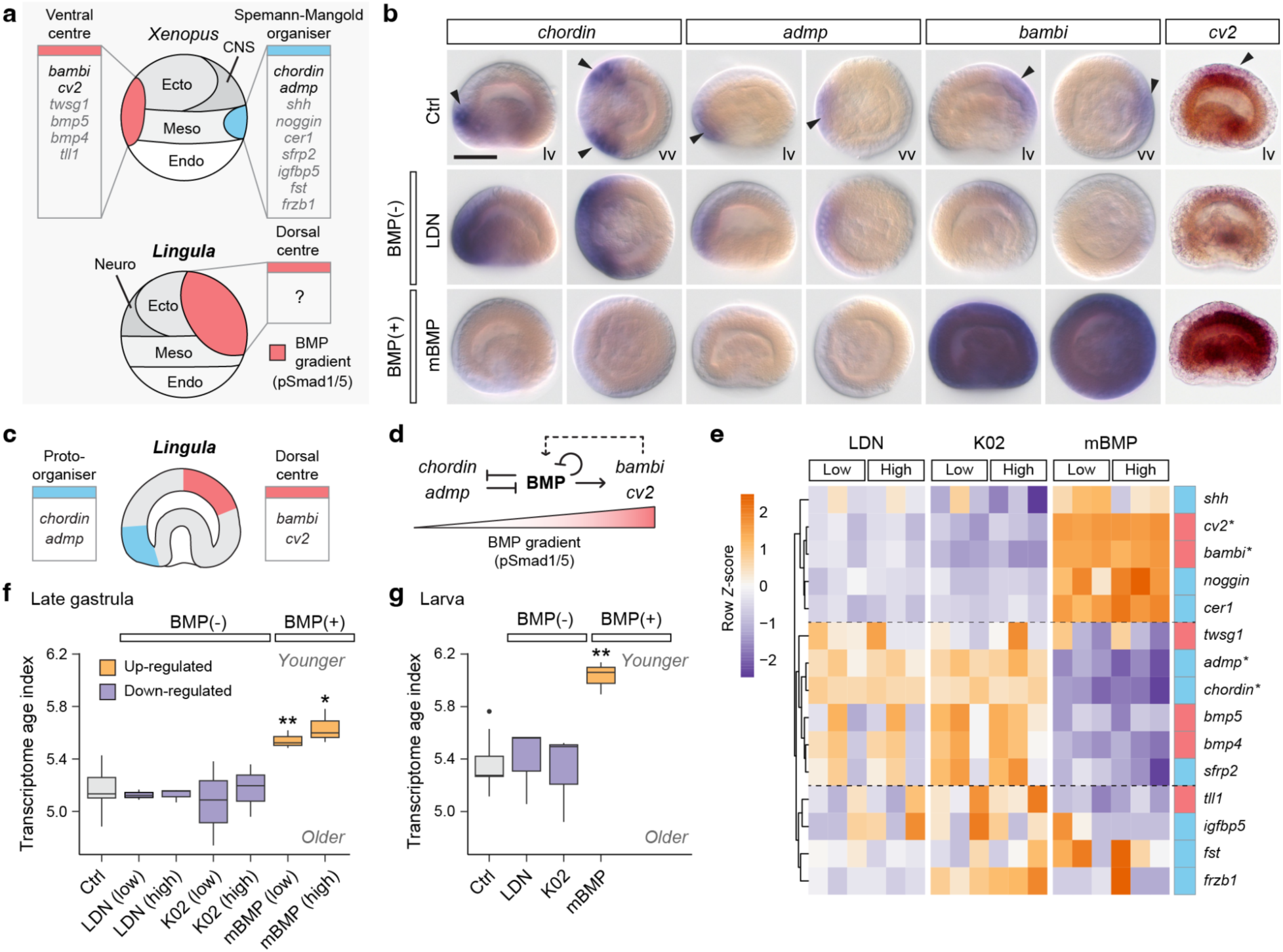
Brachiopods exhibit BMP-mediated neural patterning and a primordial organiser. **a**, Top: Schematic illustration of proteins secreted by ventral and dorsal (Spemann-Mangold organiser) signalling centres in the *Xenopus* gastrula. Bottom: Illustration of the BMP gradient in the *L. anatina* gastrula, corresponding to the ventral centre in *Xenopus*. Ecto, ectoderm; Meso, mesoderm; Endo, endoderm. **b**, The expression profile of *Xenopus* dorsal genes (*chordin* and *admp*) and ventral genes (*bambi* and *cv2*) under the manipulation of BMP signalling. lv, lateral view; vv, vegetal view. Scale bar, 50 μm. **c**, Cartoon illustration of the expression domains of kernel BMP signalling genes at the late gastrula stage. **d**, Proposed molecular network of the BMP signalling genes in a BMP– Chordin axis generating a BMP gradient (red triangle). BMP in bold indicates BMP signals. Dashed line indicates an unknown interaction between the modulators and BMP signals. **e**, Expression profile of orthologues of key *Xenopus* BMP pathway genes in the *L. anatina* gastrula under BMP signalling manipulation. Low dose (BMP, 100 ng/mL; LDN, 4 μM; K02, 250 nM); high dose (BMP, 200 ng/mL; LDN, 8 μM; K02, 500 nM). *Xenopus* ventral centre genes (red) and Spemann-Mangold organiser genes (blue) labelled on the right. Asterisks highlight genes within the *Xenopus* ventral centre and the Spemann-Mangold organiser that BMP signals similarly regulate in brachiopods. **f** and **g**, Transcriptome age index of *L. anatina* late gastrula embryos (**f**) and larvae (**g**) under conditions of BMP signal manipulation. Asterisks show significant differences from control samples at *p* < 0.05 (*) and *p* < 0.01 (**) as determined by two-sided *t*-tests. A higher transcriptome age index indicates an evolutionarily younger transcriptome.

The parallel in regulatory patterns between *Xenopus* and *L. anatina* suggests a conserved BMP-mediated dorsal–ventral patterning system architecture (Fig. 5a). We next investigated whether the spatial expression patterns of these genes in *Xenopus* are also conserved in *L. anatina*. *In situ* hybridization experiments show that *bambi* and *cv2* are dorsally localised in *L. anatina* gastrulae, while *chordin* and *admp* are found ventrally (Fig. 5b). This is consistent with their expression in *Xenopus* when the axis inversion is taken into account. Notably, both *chordin* and *admp* were expressed at the boundary of the oral ectoderm and endomesoderm, similar to the expression pattern observed in the chordate *B. floridae*^16^ (Fig. 5b). In contrast, *bambi* and *cv2* were expressed in domains with active BMP signals. BMP(-) treatment causes expansion of *chordin* and *admp* expression domains and the near absence of *bambi* and *cv2* expression, suggesting ventralisation of the gastrula. In contrast, BMP(+) treatment results in the near absence of expression of *admp* and *chordin* while *bambi* is expressed universally, suggesting gastrula dorsalisation (Fig. 5b). These experiments support our transcriptomic results showing that BMP signalling upregulates *bambi* and *cv2* while downregulating *chordin* and *admp*. This result is also consistent with regulatory interactions observed in *Xenopus*, where BMP signalling represses *chordin*^5^ and *admp*^10,79^ while activating *bambi*^77,80^ and *cv2*^5,73^. Overall, we found conservation of *Xenopus* dorsal–ventral patterning gene regulatory relationships and gene expression patterns in the brachiopod *L. anatina* (Fig. 5c,d), indicating deep conservation of these pathways in spiralians.

Despite this remarkable conservation, several *Xenopus* Spemann-Mangold organiser and ventral centre genes (e.g., *twsg*, *noggin* and *follistatin*) do not appear in our restricted differentially expressed lists (Fig. 4g,h). We hypothesised that this may reflect either our highly conservative approach to differentially expressed gene identification or genuine divergence in the role of these genes between species. To investigate this, we studied the expression of 15 key *Xenopus* Spemann-Mangold organiser and ventral centre genes under each manipulation condition. Intriguingly, many of these genes are differentially expressed in response to BMP signalling in *L. anatina*, but not in the predicted direction. For instance, Spemann-Mangold organiser genes *cer1*, *noggin* and *shh* are BMP-upregulated when they might be expected to be downregulated like *chordin* and *admp* (Fig. 5e). This finding suggests that although the central BMP–Chordin axis is conserved, its downstream effects are evolutionarily variable. Accordingly, a phylostratigraphy approach using transcriptome age index (TAI) analysis^81^ shows that increasing the level of BMP signalling significantly decreases the evolutionary age of the transcriptome during both gastrula and larval stages of development (*t*-test *p* < 0.05; Fig. 5f,g and Supplementary Tables 29 – 31). This result suggests that activating the BMP pathway in *L. anatina* leads to the upregulation of evolutionarily young genes. Indeed, our findings show that during the developmental time course, the late blastula and early gastrula stages—following BMP signal activation—exhibit the highest TAI, indicating the youngest transcriptome (Supplementary Fig. 5). This aligns with transcriptome ages at similar stages in zebrafish development^81^. This suggests that BMP signals are crucial for gastrulation, marked by the high expression of genes that emerged during bilaterian evolution and are involved in cellular interactions.

Finally, we investigated how BMP ligand expression relates to the BMP signalling gradient in *L. anatina*. *In situ* hybridisation experiments revealed that, in the ectoderm, *bmp2/4* and *bmp5-8* are expressed dorsally in the region of pSmad1/5 activation (Extended Data Fig. 9a), directly opposite *chordin* expression (Extended Data Fig. 9b). In contrast, *Bmp3* is expressed in ventral ectoderm, while all three genes show expression in the endomesoderm (Extended Data Fig. 9a). BMP manipulation experiments suggest that ectodermal *bmp2/4*, *bmp3* and *bmp-5-8* expression is autoinhibited by BMP signalling (Extended Data Fig. 9a-d). This spatial expression data suggests that dorsal Bmp2/4 localisation is a possible driver of the dorsal pSmad1/5 activation. Overall, our study shows that the *L. anatina* dorsal–ventral axis is patterned by a BMP gradient that exhibits notable similarities of that of the vertebrate *Xenopus*. This is consistent with a deeply conserved network maintained in both spiralians and deuterostomes since their split at the base of the bilaterians.

## Discussion

Elucidating the evolution of axial patterning mechanisms within bilaterians remains a key goal of evolutionary developmental biology. Until now, direct topological comparisons of early development between one of the three major clades of bilaterians and the other two groups (i.e., Spiralia with Deuterostomia and Ecdysozoa) have been prevented by the unique development of spiralians^33,34^. Our study fills this gap by using the brachiopod *L. anatina*, a spiralian with deuterostome-like development. Importantly, we find BMP signal read-out and *bmp2/4* expression at the dorsal side of the gastrula and *chordin* expression from the ventral side, consistent with ecdysozoan protostomes like *Drosophila*^11–13^ and basal deuterostomes, such as sea urchins^17,20^ and hemichordates^15,20^. Indeed, manipulations to BMP signalling in *L. anatina* interfere with proper dorsal–ventral axis patterning in both the gastrula and early larva. For instance, BMP inhibition causes the expansion of the expression domains of ventralising factors, particularly *chordin*. This has clear morphological effects including the failure of mantle folding. Conversely, BMP pathway overstimulation expands the dorsal gene expression domain and causes ‘over-folding’, accompanied by internalisation of the tentacle and mouth. Overall, these results reveal the presence of BMP-mediated dorsal–ventral patterning that, in contrast to that of other spiralians, exhibits several features that are highly conserved with distantly related bilaterians such as *Drosophila* and *Xenopus*.

The similarities between gene expression localisation and regulatory interactions in *L. anatina* and *Xenopus*, particularly those regarding the Spemann-Mangold organiser, are especially striking. In the basal chordate amphioxus, such similarities have been taken, along with transplantation experiments^82^, as evidence that the organiser was present at the base of the chordates^16,83^ or the deuterostomes^84^, not just in vertebrates. In light of the extension of these similarities to *L. anatina*, we consider it a possibility that these features have been inherited and maintained in both lineages from a bilaterian ancestor that possessed a ‘proto-organiser’. Transplantation experiments testing whether part of the dorsal region of *L. anatina* embryos possesses organiser capabilities should therefore be a priority, although in any case homology would be difficult to assess. Irrespective of the presence or absence of an organiser in *L. anatina*, the results of this study suggest deep evolutionary conservation between brachiopods and deuterostomes that supports the hypothesis that a classical Bmp2/4–Chordin dorsal–ventral patterning system was present in the ancestor of spiralians despite their derived mechanism of early embryonic development (Fig. 6).

**Fig. 6.**
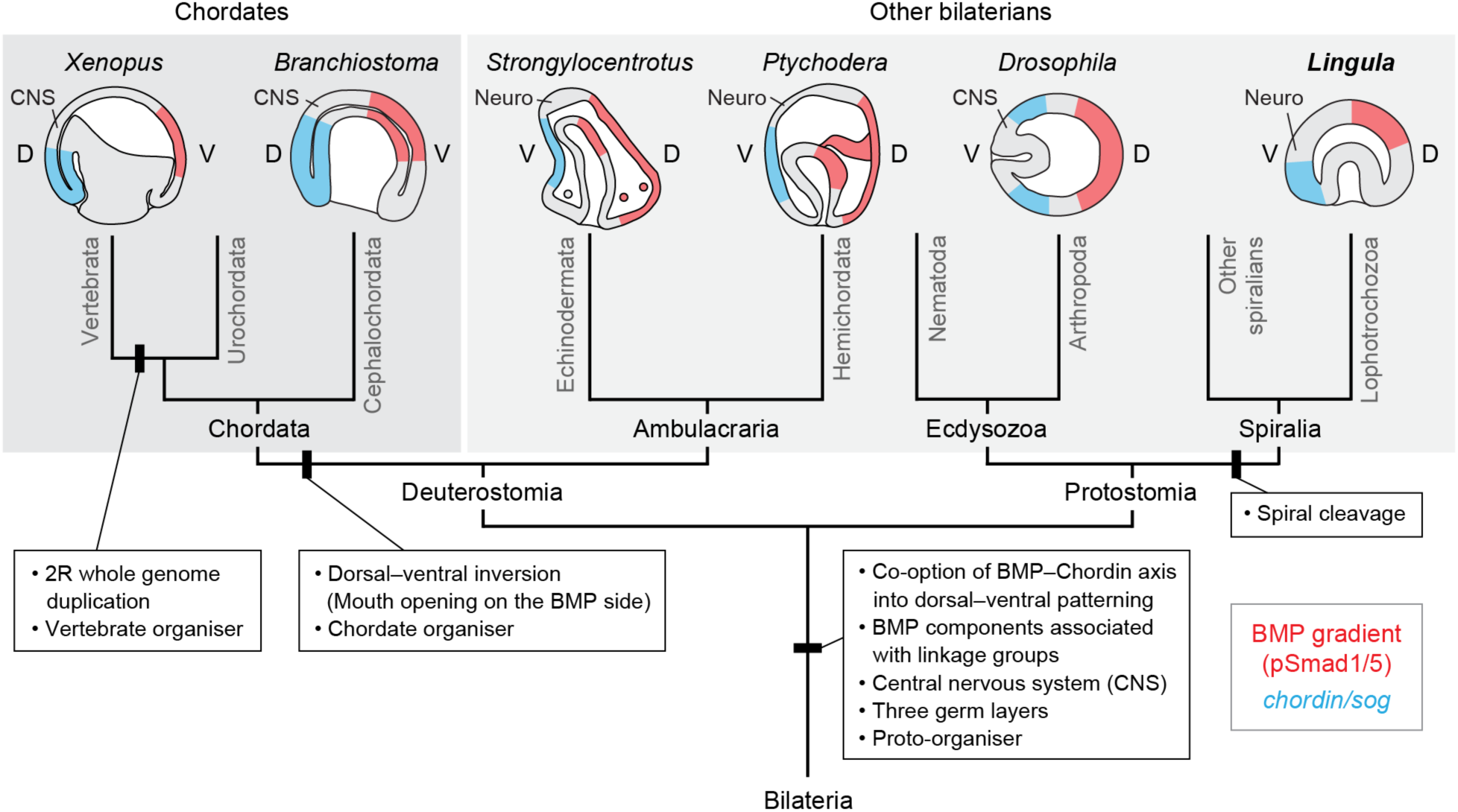
Evolution of BMP gradients and dorsal–ventral patterning in bilaterians. Although the spatial expression patterns of BMP ligands are variable, the domain with high BMP signalling readout (pSmad1/5, red) and the anti-BMP domain (*chordin*/*sog*, blue) are on the dorsal and ventral sides, respectively, of the ecdysozoan and basal deuterostome embryos. Our data from *L. anatina* shows that this is also the case in spiralians. Due to dorsal-ventral axis inversion, this is reversed in chordates. However, the core molecular patterning mechanisms remain unchanged, especially for those of kernel modulators and antagonists. In all cases, the neural domain develops on the side at which anti-BMP signals are produced, since BMP signalling inhibits neural (Neuro) and central nervous system (CNS) formation in most bilaterians.

One significant observation casting doubt on the ancestral state of BMP-mediated dorsal–ventral axis patterning in spiralians^27^ was the discovery that, in some molluscs and annelids, BMP signals have no effect or even a positive effect on neural development^24,31,32^. This stands completely contrary to the predicted effect given results in ecdysozoans and deuterostomes^21^. Through functional transcriptomics of embryos with manipulated levels of BMP signalling, we find strong inhibition of neural genes by the *L. anatina* BMP pathway during development. This result demonstrates that the function of BMP signalling in restricting neural induction is also conserved in some spiralians and supports the contention that this was the ancestral state of Spiralia.

Although core features of the dorsal–ventral patterning BMP pathway (e.g., *chordin*, *admp*, *bambi* and *cv2*) are conserved from *Xenopus* to brachiopods, many genes with important roles in *Xenopus*, such as *twsg*^74–76^, *noggin*^8,9^ and *follistatin*^78^, are not regulated by BMP signaling in the same way in *L. anatina*. This finding suggests that while the central BMP– Chordin dorsal–ventral patterning mechanism is highly conserved, peripheral components of the gene regulatory network show considerably more evolutionary flexibility. Consistent with this, we also find that the *L. anatina* BMP pathway upregulates the expression of evolutionarily young genes. Variations in the role of BMP signalling in dorsal–ventral axis development in spiralians, such as the positive effect on neurogenesis in some molluscs, may therefore arise from lineage-specific divergence in the pathway’s downstream responses, rather than a different ancestral role in spiralians. Our result supports a model where the core function of the BMP pathway itself is ancient and deeply conserved but the downstream effects involve evolutionarily younger and lineage-specific genes.

## Methods

### Biological materials and artificial fertilisation

Between 2013 to 2016, gravid adults of the brachiopod *L. anatina* (approximately 150 to 200) were collected each August during low tide from Kasari Bay, Amami Island, Japan (28.440583 N, 129.667608 E). Around 50 individuals were housed in a 10-L aerated seawater tank and fed daily with 25 mL/tank of *Chaetoceros calcitrans* (5×10^7^ cells/ml). Oocyte maturation was induced by injecting each gonad with 30 μl of 40 mM dibutyryl-cAMP dissolved in phosphate-buffered saline (PBS)^37,42^. To control fertilisation timing, post-injection individuals were isolated in separate Petri dishes. Artificial spawning was facilitated by adjusting the temperature from 25°C to 29°C over 2 h and then rapidly returning it to 25°C for a cold shock^85^. Prior to inducing fertilisation, sperm activity was inspected under a stereomicroscope. Fertilisation efficacy decreased sharply 4 h after spawning. Depending on seasonal factors, spawning rates varied from 20% to 70%.

### Genomic sample acquisition and DNA extraction

In August 2021, fresh adults of *L. anatina* were collected with the objective of obtaining high-quality genomic DNA suitable for both long-read and Hi-C sequencing methodologies. The dissected tissues, including the mantle, lophophore, gonad, adductor muscle, and pedicle, were immediately snap-frozen using liquid nitrogen. For genome resequencing, approximately 2 g of mantle and lophophore tissues and 1.5 g of adductor muscle tissue were collected. These tissues were then divided into two equal portions. One portion was used for genomic DNA extraction, while the other was reserved for Hi-C sequencing. The cetrimonium bromide (CTAB) method was utilised to extract high molecular weight genomic DNA. After extraction, the DNA was purified using the Blood & Cell Culture DNA Midi Kit (Qiagen 13343). DNA purity was assessed with a NanoDrop spectrophotometer, and DNA concentration was determined using the Qubit 4.0 Fluorometer.

### RNA sequencing of adult tissues

Total RNA was extracted from the mantle, gonad, lophophore, adductor muscle and pedicle using TRIzol. The RNA was reconstituted in 50 μl RNase-free water, and its concentration was determined with a Nanodrop spectrophotometer. RNA quality was verified using the 5400 Fragment Analyzer System (Agilent Technologies). cDNA libraries for each tissue sample were constructed with the TruSeq RNA Library Prep Kit v2. Libraries were sequenced as 150 bp paired-end reads on the Illumina NovaSeq 6000. Quality control was conducted using FastQC^86^. The reads were trimmed with Trimmomatic (v0.39)^87^ and then aligned to the genome using STAR (v2.7.10b)^88^ for gene prediction (Supplementary Table 32). For gene expression analysis, transcript abundances were quantified using kallisto (v0.43.0)^89^. The same method was used to process data from a published developmental dataset of *L. anatina*^37^ (PRJNA286275) (Supplementary Table 33).

### Genome sequencing and assembly

For PacBio HiFi circular consensus sequencing, SMRTbell libraries were constructed following PacBio’s standard protocol, utilising the 15-kb preparation solutions. In brief, 15 μg of DNA was extracted from the mantle, lophophore and adductor muscle tissue samples (5 μg from each tissue) and was used for DNA library preparations. The genomic DNA was sheared to the desired fragment size using g-TUBEs (Covaris). After removal of single-strand overhangs, damage repair, end-repair and A-tailing, DNA fragments were ligated with the hairpin adaptor. Post-ligation, the library was treated with nuclease, cleaned using the SMRTbell Enzyme Cleanup Kit, and purified with AMPure PB Beads (Beckman Coulter). The desired fragments were subsequently isolated using BluePippin (Sage Science). An Agilent 2100 Bioanalyzer was used to determine the size distribution of the library fragments. The final sequencing of the SMRTbell library was conducted on the PacBio Sequel II platform, utilising Sequencing Primer V2 and the Sequel II Binding 2.0 Kit at Grandomics Biosciences. The Hi-C library was constructed following a previously described method^90^. In brief, mantle, lophophore, and adductor muscle tissues were cut into 2 cm segments and cross-linked using a nuclei isolation buffer with 2% formaldehyde. Post cross-linking, tissues were ground to produce a nuclei suspension. The isolated nuclei were digested with 100 units of DpnII and labelled with biotin-14-dATP. Unligated fragments had their biotin removed using T4 DNA polymerase. The DNA was sheared to 300–600 bp fragments, underwent blunt-end repair and A-tailing, and was isolated with streptavidin beads. After quality checks with a Qubit Fluorometer and an Agilent 2100 Bioanalyzer, the libraries were sequenced as 150 bp paired-end reads on the Illumina NovaSeq 6000 platform.

Hi-C data was used for assembly scaffolding^91^ using the following pipeline: trimmomatic (quality filtering)^87^; FastQC (quality control)^86^; bwa_mem2 (sequence alignment)^92^; Juicer (data filtering)^93^; 3D-DNA (Hi-C-assisted assembly)^94^; and JuiceBox (visualisation and error correction)^95^. Having obtained a preliminary assembly, blasts, read coverage and GC content were used to check for sequences belonging to contaminants and symbionts, The assembly was first subjected to Diamond^96^ blastx against the nr database of NCBI with ‘sensitive mode’ parameter and an E-value of 1e-25. Subsequently, Hi-C reads were mapped to the original genome using HISAT2^97^ with default parameters and in single-end mode to calculate the coverage of each fragment. These three data types were visualised using blob plots created with BlobToolKit^98^. This methodology revealed a small cluster of short sequences with an unusually high GC content (greater than 45%) which blasted to non-brachiopod taxa. All such fragments were removed from the genome. A second blob plot, produced from the remaining fragments, identified only sequences blasting to Brachiopoda. Sequences present after this filtering process represent the final *L. anatina* assembly.

### Gene prediction and annotation

Repetitive elements were annotated *de novo* using RepeatModeler (v2.0.4)^99^. RepeatModeler employs RepeatScout^100^ and RECON^101^ to identify transposable elements and also uses the long terminal repeat (LTR)-specific tools LTRharvest^102^ and LTR_retriever^103^. The BRAKER pipeline (v3.0.3)^104–112^ was then used for gene prediction and annotation. Genomes were first soft-masked with RepeatMasker (v4.1.5; sensitive mode)^113^, and then BRAKER was run using hints from mapped RNA sequencing data. RNA-seq reads (Supplementary Table 34) were downloaded from the NCBI Sequence Read Archive, trimmed with Trimmomatic (v0.39)^87^ and aligned with STAR (v2.7.10b)^88^ before input to BRAKER in BAM format. The RNA-seq datasets generated in this study were also used as hints for BRAKER. Gene annotation quality was assessed using BUSCO (v5.4.7)^114^. InterProScan^115^, KofamScan^116^ and EggNOG-mappe^117,118^ were used for functional annotation. For KofamScan, output is limited to hits where threshold > score (adjudged to be a significant hit; see^119^ for complete explanation). Orthologues of *L. anatina* genes in human and mollusc (*Patella vulgata*) genomes were identified with OrthoFinder^120^. RepeatLandscape^113^ was used to create Kimura substitution level plots for repeats in lophophorate genomes. Ribosomal RNA (rRNA) genes were annotated with barrnap (v0.9)^121^. Gene density and repeat density plots were made with RIdeogram (v0.2.2)^122^.

### Phylogenetic analysis

Proteomes were downloaded from NCBI or produced by the gene prediction method outlined above (Supplementary Table 12). Orthology assignment was performed with OrthoFinder (v2.5.4)^120^. OrthoSNAP (v0.01)^123^ was then run with default parameters to recover additional orthologues for phylogenetics. The resultant dataset contained 2,036 OrthoSNAP orthogroups. To determine the strength of the phylogenetic signal possessed by each orthogroup, we first aligned sequences with MAFFT (v7.520)^124,125^ and trimmed alignments with ClipKIT (v1.4.1)^126^. Individual gene trees were then constructed with IQ-TREE (v2.2.2.3)^127^ using ModelFinder automated model selection^128^ and UFBoot2 ultra-fast bootstrapping^129^. Orthogroups with an average bootstrap score over 85 % (*n* = 109) were selected for species tree building and alignments concatenated with PhyKIT (v1.11.7)^130^. The final tree was built using IQ-TREE as above (model LG+F+R5). In order to assess whether constructing trees using orthologues with lower mean bootstrap scores (weaker phylogenetic signal) gives different results, orthologues were subsetted by mean bootstrap scores at 5% intervals, and trees were constructed for each group as above.

### Comparative genomics

We used OrthoFinder (v2.5.4)^120^ to identify BMP-related components in lophotrochozoan genomes. All cases of putative gene losses and duplications were manually verified using approaches such as reciprocal blast searches, microsynteny comparisons and gene tree construction with IQ-TREE (v2.2.2.3)^127^. Attempts to assess the functionality of genes based solely on genomic sequence can lead to the erroneous designation of functional genes as pseudogenes. Thus, for the purpose of this analysis, we consider genes as only the number of copies of a gene and do not speculate on functionality. Human BMP-related genes are well-characterised and gene counts are not re-assessed here.

Gene family evolution was modelled using CAFE (v5.0.0)^131^. CAFE implements a stochastic birth and death model to estimate the number of gene gains and losses occurring at each node in a tree. The species set used for this analysis was identical to that used for phylogeny reconstruction with one exception: the nemertean *Notospermus geniculatus* was removed because the published genome has high levels of redundancy, which would result in inaccurate estimates. To run CAFE, an ultrametric version of the above species tree was calculated using pyr8s from iTaxoTools^132^ to implement r8s^133^. OrthoFinder was used as above for the orthology assignment. Principal component analysis (PCA) was performed on the orthogroup-species matrix using R.

### Homology modelling

Homology modelling of the *L. anatina Smad1/5* protein was completed using Modeller^134^ in UCSF Chimera (v1.15)^135^. The human SMAD1 protein was used as a reference.

### Macrosynteny analysis

Macrosynteny was compared between *L. anatina* and species representing four other phyla with relatively conserved genomic organisations: *B. floridae* (Florida lancelet, Chordata), *P. maximus* (scallop, Mollusca), *L. longissimus* (bootlace worm, Nemertea) and *O. fusiformis* (Annelida). Proteomes for these species were obtained from NCBI. OrthoFinder (v2.5.4)^120^ was then used to identify single-copy orthologues of *B. floridae* genes assigned to a bilaterian ancestral linkage group (ALG) by Simakov et al. (2022)^41^. SyntenyFinder^136^, which implements the R package RIdeogram (v0.2.2)^122^, was used to create ideogram plots and Oxford dot plots. The genomic locations of BMP pathway-related genes, Wnt ligands, and Hox genes in each of the five species were identified with blast (v2.14.1). Conserved associations with ALGs were inferred from the genes’ chromosomal locations. The rate of conserved associations of developmental genes with a specific ALG was compared to that of a random sample of 100 single-copy orthologues using a chi-square test (Supplementary Table 35).

### Manipulation of BMP signals

BMP signalling manipulation experiments were deployed to reveal the function of the BMP pathway during *L. anatina* development. Two separate small molecule inhibitors were used to block the BMP pathway: LDN193189 (LDN; Stemgent 04-0074)^59^ and K02288 (K02; Tocris 4986)^60^. These conditions are referred to as BMP(-). Exogenous mouse recombinant BMP4 (mBMP4; R&D Systems 5020-BP) was used to over-activate the BMP pathway^61^. This condition is referred to as BMP(+). Manipulations were applied for two different durations, from the early blastula stage (5 h post fertilisation, hpf) to either the late gastrula (10 hpf) or the 1-pair-cirri larval stage (24–27 hpf). Two doses (high and low) of each manipulator were applied to the late gastrula experiments: LDN (4 and 8 μM), K02 (250 and 500 nM) and mBMP4 (100 and 200 ng/mL). One dose was applied to the 1-pair-cirri larva experiments: LDN (2 μM), K02 (250 nM) and mBMP4 (200 ng/mL). Controls were run for each manipulator with the vehicle only (bovine serum albumin (BSA) or dimethyl sulfoxide).

### Functional transcriptomics

RNA sequencing was used to explore the impacts of BMP signal manipulation experiments on the transcriptome. RNA was extracted for three biological replicates of each condition (45 samples, Supplementary Table 21) using TRIzol and sequenced using the Illumina HiSeq 4000 platform (total 855 million read pairs). After performing quality control with FastQC (v0.11.5) and trimming with Trimmomatic (v0.36), transcript abundances were quantified using kallisto (v0.43.0)^89^. Differential expression analysis was conducted using a bundled script in Trinity (v2.3.2)^137^, primarily utilising edgeR with a dispersion parameter of 0.1^138^. Transcripts were considered statistically differentially expressed with a false discovery rate (FDR) of less than 0.05. Pearson’s correlation coefficient was used to assess the similarity of each condition’s transcriptome. Putative BMP signalling downstream genes were identified based on their expression changes (at least fold-change > 2, *p* < 0.01). Genes that were upregulated in the BMP(+) condition and downregulated in the BMP(-) condition were classified as BMP-upregulated. Conversely, genes that were downregulated in the BMP(+) condition and upregulated in the BMP(-) condition were classified as BMP-downregulated. Gene ontology analysis was performed on upregulated and downregulated gene sets using reciprocal best hits from BLAST searches against the Swiss-Prot database from UniProt^139^. To focus on the most robustly differentially expressed genes, we limited the dataset to genes that were differentially expressed in both the late gastrula and 1-pair-cirri-larva experiments. Differentially expressed gene sets were searched for genes known to be involved in dorsal– ventral patterning in *Xenopus*.

### Transcriptome age index analysis

The phylostratigraphic age of each gene in the *L. anatina* genome was first estimated using GenEra (v1.2.0)^140,141^. GenEra reduces biases in age assignment by using DIAMOND^96^ to search the entire NR database (Supplementary Table 36). To improve the resolution of age assignment in the case of *L. anatina*, we also added one genome assembly for each animal phylum that does not have a RefSeq annotated genome (Supplementary Table 37). This is especially important within the Spiralia, where key phyla like Bryozoa and Phoronida are unrepresented. Using the estimated gene ages, the R package myTAI (v1.0.1.9000)^142^ was used to calculate the transcriptomic age index^81^ for several *L. anatina* transcriptomic datasets.

These were (i) a developmental time course from fertilised egg to two pair cirri larva (ii) *L. anatina* adult tissues (adductor muscle, lophophore, ovary, mantle and pedicle) and (iii) *L. anatina* late gastrula and larval stages with BMP signalling manipulation.

### Bacterial-cloning-free riboprobe preparation

To prepare DNA templates for RNA probe synthesis, a bacterial-cloning-free protocol was developed to maintain both rapid preparation and target specificity (Supplementary Fig. 6). This protocol initially uses gene-specific primers (F1 and R1, designed from transcriptomes) to amplify target sequences in the first PCR. The amplified sequences are then ligated into the pGEM-T Easy Vector to attach an RNA polymerase promoter site. Inserts in the reverse direction (antisense) to the T7 promoter site were further amplified using T7 and gene-specific forward nested primers (F2) in a second PCR. This nested PCR ensures both the specificity of the PCR products and the correct transcriptional direction of the inserts. The products from the second PCR can then be used for sequencing validation and *in vitro* transcription. No cloning-based screening is required to select target clones, allowing the entire procedure to be completed within one day.

Total RNA from various embryonic stages was extracted using TRIzol and cleaned up with the RNeasy Micro Kit (Qiagen 74004). cDNA synthesis was performed using the SuperScript III First-Strand Synthesis System (Thermo Fisher Scientific 18080051). Target sequences were amplified in a first PCR using gene-specific primers (F1 and R1) and EmeraldAmp GT PCR Master Mix (TaKaRa RR320A). The PCR conditions were: 94°C for 2 min, followed by 30 cycles of 94°C for 15 seconds, 55°C for 15 seconds, and 72°C for 1 min 30 seconds, with a final extension at 72°C for 2 min. PCR products were analysed on a 1% agarose TAE gel stained with SYBR Safe DNA Gel Stain (Thermo Fisher Scientific S33102), and remaining products were purified using the QIAquick PCR Purification Kit (Qiagen 28104). Purified PCR products were ligated into the pGEM-T Easy Vector (Promega A1360) using 5 μl of 2X Rapid Ligation Buffer, 0.5 μl of pGEM-T Easy Vector (25 ng), 3.5 μl of PCR product, and 1 μl of T4 DNA Ligase in a 10 μl reaction, incubated for 1 h at room temperature. A second PCR was performed using T7 and gene-specific forward nested primers (F2). The PCR conditions were: 94°C for 2 min, followed by 35 cycles of 94°C for 15 seconds, 55°C for 15 seconds, and 72°C for 1 min 30 seconds, with a final extension at 72°C for 2 min. PCR products were analysed on a 1% agarose TAE gel, and target bands were purified using the Wizard SV Gel and PCR Clean-Up System (Promega A9282).

*In vitro* transcription was performed using T7 RNA polymerase (Promega RP2075) and DIG RNA Labeling Mix (Roche 11277073910). The reaction was incubated for 2–4 h at 37°C, followed by DNase I treatment for 15 min at 37°C. RNA probes were precipitated with 50 μl of nuclease-free water, 30 μl of 7.5 M LiCl, and 300 μl of 100% ethanol, chilled at -20°C for at least 30 min, and centrifuged at 16,000x*g* for 20 min at 4°C. The RNA pellet was washed with 300 μl of cold 80% ethanol, air-dried, and resuspended in 25 μl of nuclease-free water. RNA probes were then diluted with 25 μl of 100% formamide and stored at -20°C or -80°C.

### *In situ* hybridisation for localising gene expression

Embryos were fixed overnight at 4°C with 4% paraformaldehyde (PFA; Electron Microscopy Sciences 15714) in filtered seawater. Post-fixation, embryos were washed with filtered seawater, dehydrated in 100% methanol, and stored at -20°C. For rehydration, embryos were transferred from methanol into baskets with 40-μm nylon mesh, immersed in PBST (0.1% Tween 20 in PBS) in a 24-well plate, and incubated for 10 min with 1 ml per well. Permeabilisation was performed in PBSN (1% NP-40 and 1% SDS in PBS) for 10 min, followed by PBSTX (0.2% Triton X-100 in PBS) for 10 min. Optional bleaching was done in 2% H2O2 in PBST at room temperature for 30–60 min under direct light. Embryos were washed in PBST for 5 min, repeated three times, and rinsed with wash buffer (50% Formamide, 5X SSC, 1% SDS, 5 mM EDTA, and 0.1% Tween 20). Prehybridization was conducted in hybridization buffer (50% Formamide, 5% Dextran sulfate, 5X SSC, 1% SDS, 1X Denhardt’s, 100 μg/ml Torula RNA, 5 mM EDTA, 0.1% Tween 20, and 50 μg/ml Heparin) at 60°C for at least 1 h with slight rocking, with plates sealed in a plastic bag or covered with plastic wrap to prevent evaporation.

For hybridisation, the hybridisation buffer with probe (1:100–1:250, >100 ng/ml) was preheated at 70°C for 5 min, then applied to the embryos and incubated at 60°C overnight with slight rocking. Post-hybridisation washes were done at 60°C for 15 min each, with three washes in wash buffer, followed by washes with wash buffer + 2X SSCT (1:1), 2X SSCT, 0.2X SSCT, and 0.1X SSCT. Embryos were then rinsed with wash buffer at room temperature for 15 min per wash. Blocking was performed in MAB blocking buffer (100 mM Maleic acid, 150 mM NaCl, 2% Blocking reagent [Roche, 1096176], 10% Sheep serum, 0.1% Tween 20, and 0.2% Triton X-100) at room temperature for at least 1 h. Primary antibody incubation was carried out overnight at 4°C using anti-DIG-AP solution (Roche 11093274910) in MAB blocking buffer. Embryos were washed in MABTX (100 mM Maleic acid, 150 mM NaCl with 0.1% Tween 20 and 0.2% Triton X-100) five times for 20 min each, followed by two washes in TMN buffer (100 mM NaCl, 50 mM MgCl2, 100 mM Tris-Cl, 0.05% Tween 20) for 5 min each. For the chromogenic reaction, embryos were transferred to clean wells to avoid staining from impurities. The reaction was carried out using BM Purple (Roche 1442074) in the dark without rocking, ranging from 20 min to several days depending on the probe. The reaction was stopped by washing in PBST, fixing in 4% PFA/PBST for 20–30 min, washing in 100% ethanol for 5–10 min, and rinsing in PBST. Finally, embryos were mounted in 70% glycerol in PBS with 0.1% NaN3 and stored overnight for full immersion at 4°C, protected from light.

*In situ* hybridisation of *cv2*, *bmp2/4* and *bmp5–8* was performed using traditional cloning methods and visualised with NBT/BCIP stock solution (Roche 11681451001), as previously described^143^.

### Immunostaining and imaging

Immunostaining was performed to identify the presence of phospho-Smad1/5 (pSmad1/5; BMP signalling readout) and phospho-Histone H3 (pHistone H3, indicator of mitotic cell at metaphase) during *L. anatina* development under control and BMP signal manipulation conditions. Embryos from early cleavage to larval stages were fixed in 4% PFA in filtered seawater, followed by dehydration in chilled methanol and storage at -20°C. For immunostaining, embryos were first rehydrated with PBST for 10 min and then subjected to permeabilisation using PBSTX for 30 min. To block non-specific antigens, embryos were treated with 3% BSA in PBST for at least 1 h. They were subsequently incubated with either rabbit anti-phospho-Smad1/5 (1:200; Cell signalling 9511S) or rabbit anti-phospho-Histone H3 (Ser10) (1:100; Millipore 06-570) antibody in 3% BSA in PBST overnight at 4°C. Alexa Fluor goat anti-rabbit secondary antibody (1:400; Invitrogen A-11037) was used for signal visualisation of the primary antibodies. For chitin detection, a fluorescein-conjugated chitin-binding probe (1:200; NEB P5211S) was applied. Nuclei were stained with Hoechst 33342 (1:1000 dilution from a 10 mg/mL solution; Invitrogen H1399), and cytoplasmic membranes were labelled using CellMask Deep Red (1:2000; Invitrogen C10046). Imaging was performed on a Zeiss LSM 710 or LSM 780 confocal microscope.

## Data availability

Genome sequencing and RNA-sequencing datasets are accessible through NCBI BioProject (PRJNA1068743).

## Code availability

Code for analyses in this study is available at https://github.com/symgenoevolab/lingula_genome.

## Acknowledgments

We acknowledge the permission from Amami City, Kagoshima, Japan, for collecting *L. anatina* specimens. We thank Ryo Koyanagi in OIST DNA Sequencing Section for his support of RNA sequencing. We thank Asuka Sentoku for her support with the embryonic experiment at the University of Ryukyu. We also thank Keisuke Nakashima for a gift of chitin-binding probes. This work was funded by a Japan Society for the Promotion of Science Grant-in-Aid for JSPS Fellows (15J01101), a Royal Society Newton International Fellowship (NIF\R1\201315) and an Academia Sinica Career Development Award (AS-CDA-112-L06) to Y.-J.L. Y.H.W. was supported by the Innovation Team Project of Universities in Guangdong Province (2023KCXTD028) and the National Natural Science Foundation of China General Program (42276104). We thank the Symbiosis Genomics & Evolution Lab members for their assistance and support.

## Author contributions

T.D.L., Y.H.W. and Y.-J.L. conceived the project. K.S., K.E., N.S. and Y.-J.L. collected specimens. Y.H.W. sequenced the chromosome-scale genome and tissue transcriptomes.

T.D.L. and M.-E.C. annotated the genomes. K.S. and Y.-J.L. conducted embryonic experiments and functional transcriptomics. T.D.L., I.J.-Y.L. and Y.-J.L. prepared code for GitHub. T.D.L. and Y.-J.L. analysed data and wrote the manuscript. K.E., N.S., P.W.H.H. and Y.H.W. discussed the results and edited the manuscript. All authors contributed to the revision of the manuscript.

## Extended Data Figures

**Fig. S1.**
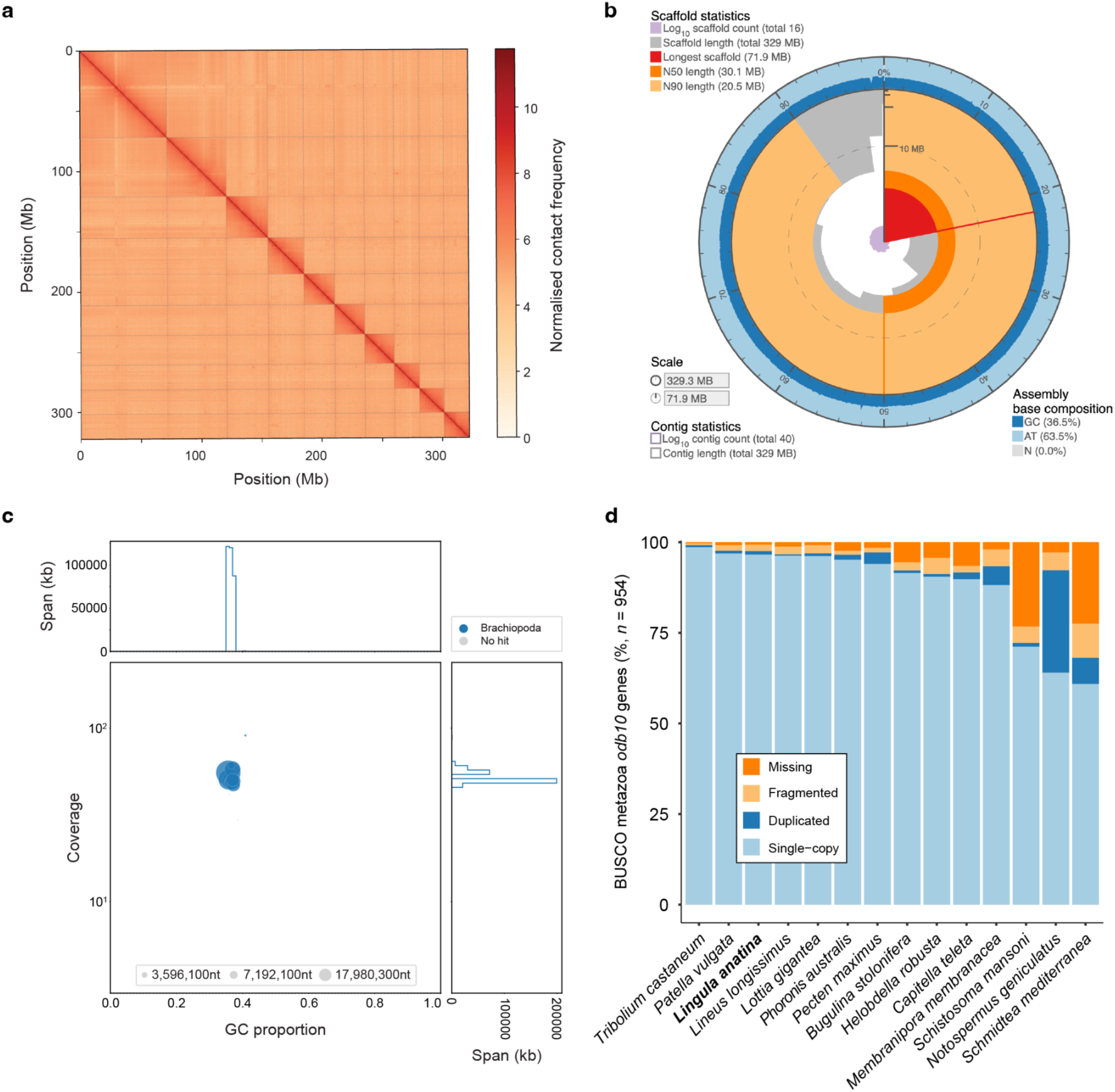
A chromosome-level assembly of the *L. anatina* genome. **a**, Hi-C contact map of the *L. anatina* genome assembly. Axes are sorted by chromosome size with highest at the top left. Colours represent intensity of interaction: darker colour = stronger interaction. **b**, Snail plot showing key statistics for *L. anatina* genome assembly. The 329.3 Mb assembly is divided into 16 scaffolds, 10 of which are chromosome-scale. The longest scaffold (red) is 71.9 Mb, the scaffold N50 (bright orange) is 30.1 Mb and the N90 (pale orange) is 20.5 Mb. The genome has a 36.5% GC content. **c**, Blob plot for the final *L. anatina* assembly showing GC content, coverage, scaffold length and blast hits. **d**, BUSCO metazoa odb10 results for selected spiralian genomes (*n* = 954). Statistics for *L. anatina*: Complete: 97.5% [Single-copy: 96.6%, Duplicated: 0.9%], Fragmented: 1.8%, Missing: 0.7%.

**Fig. S2.**
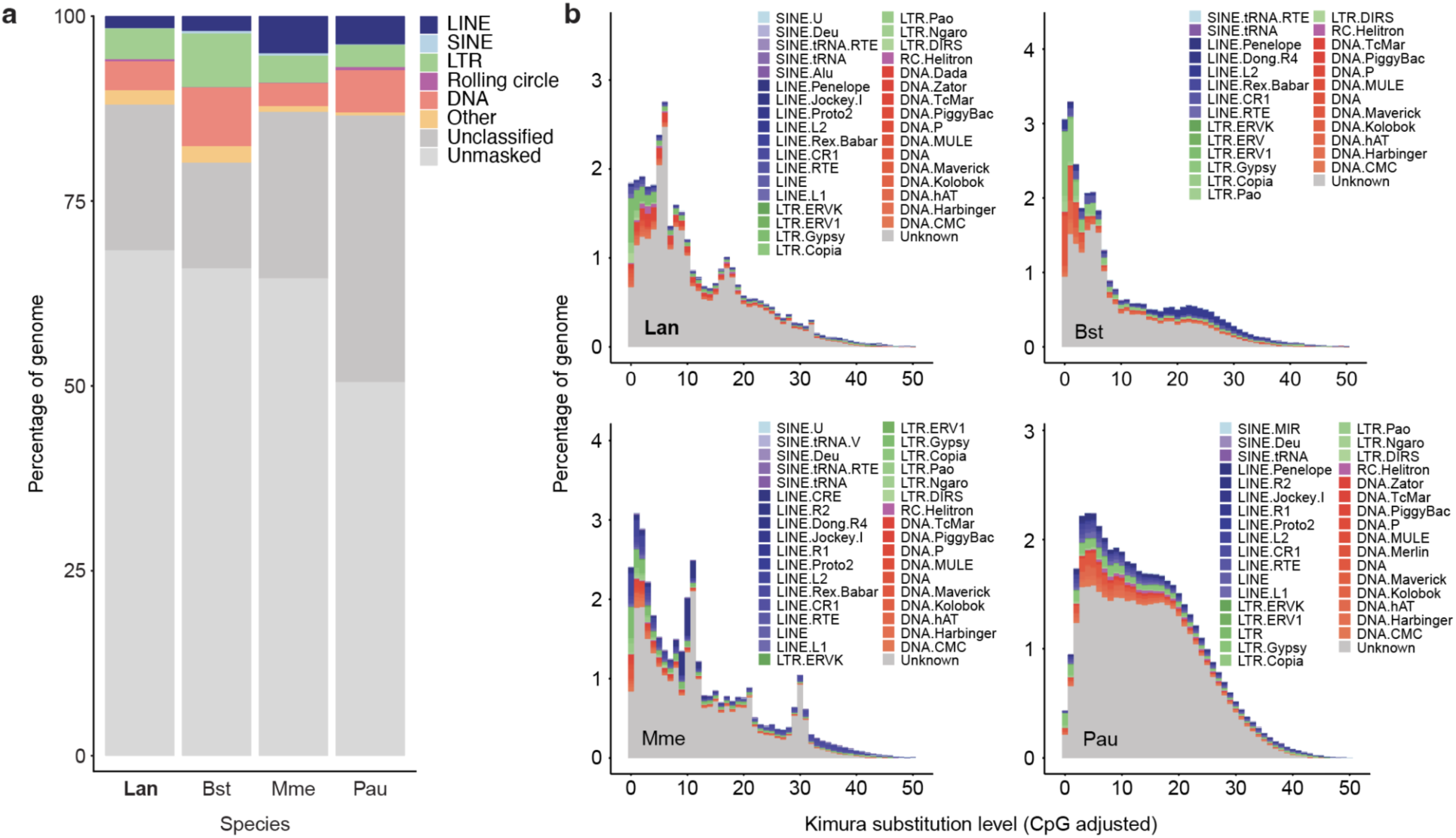
Repeat content of Lophophorata genomes. **a**, Percent of genome assemblies composed of each repetitive element group for *Lingula anatina*, *Phoronis australis*, *Bugulina stolonifera* and *Membranipora membranacea* genomes. ‘Other’ includes Penelope elements, satellites, simple repeats and low complexity elements. *L. anatina* and the two bryozoans (*B. stolonifera* and *M. membranacea*) have highly similar repeat content composition. **b**, Output from RepeatLandscape. Plots show Kimura substitution level against percent of genome occupied and are coloured by transposable element taxonomy. Kimura substitution level is a proxy for transposable element age; the higher the substitution, the older the element insertion. There are recent repeat expansions in all Lophophorata genomes. In *L. anatina* and both bryozoans, a large proportion of the annotated elements with recent expansions are long terminal repeats (LTRs).

**Fig. S3.**
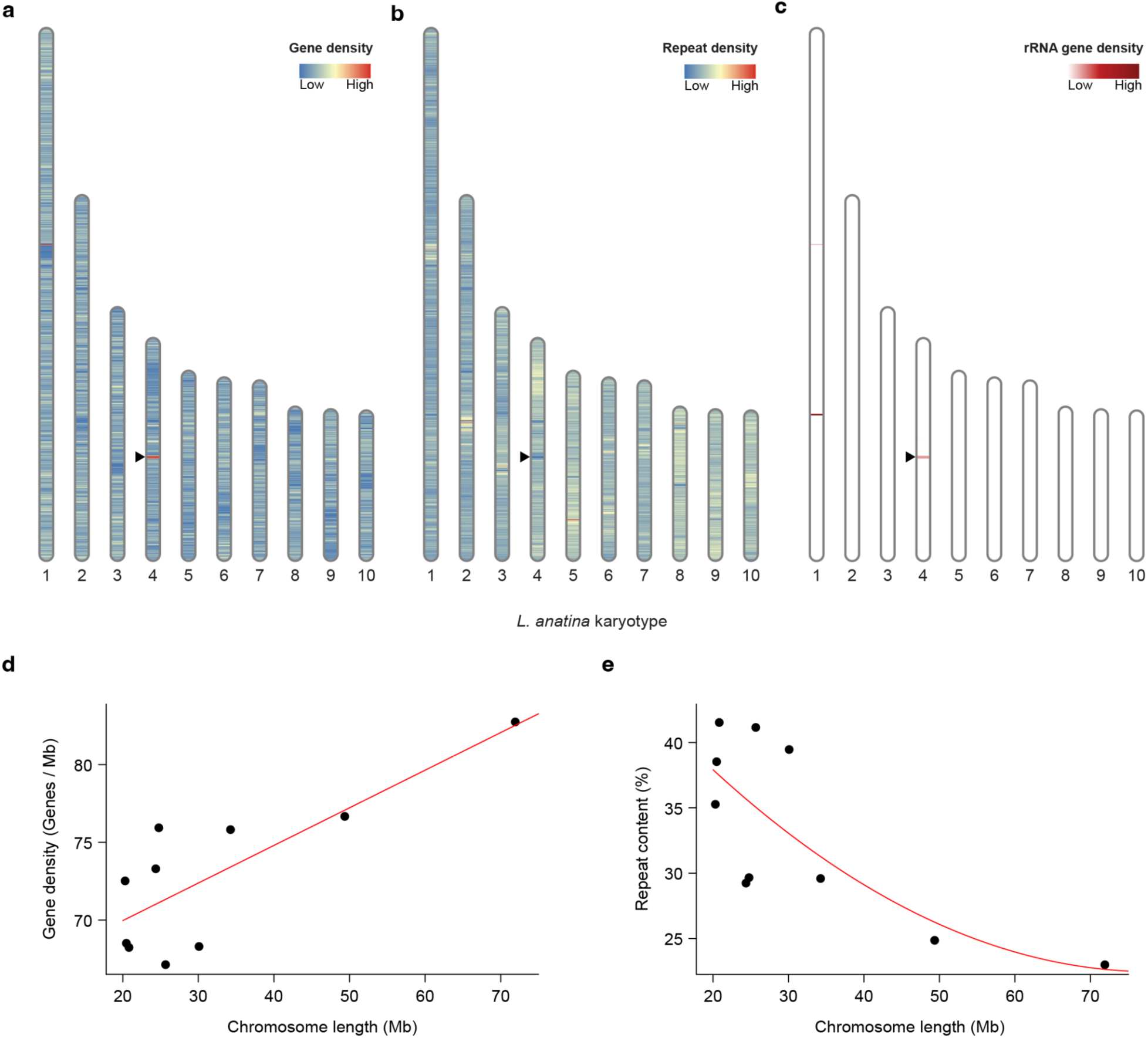
Feature density in the *L. anatina* genome. Vertical bars represent chromosomes. Colours reflect feature density. Density plots were created with RIdeogram using windows of 100 kb. Black arrowheads mark a region on chromosome 4 with exceptionally high gene density and low repeat density; this region is a cluster of ribosomal RNA (rRNA) genes. **a**, Gene density in the *L. anatina* genome. **b**, Repeat density in the *L. anatina* genome. **c**, rRNA gene density in the *L. anatina* genome. The genomic region on chromosome 4 with high gene density and low repeat density is an rRNA gene cluster. **d**, Gene density plotted against chromosome length in *L. anatina*. Longer chromosomes have higher gene density (R-squared = 0.593, F-statistic = 14.120, *p*-value = 0.006). **e**, Repeat content plotted against chromosome length in *L. anatina*. Longer chromosomes have a non-significant lower percentage of repeats (R-squared = 0.441, F-statistic = 4.556, *p*-value = 0.054).

**Fig. S4.**
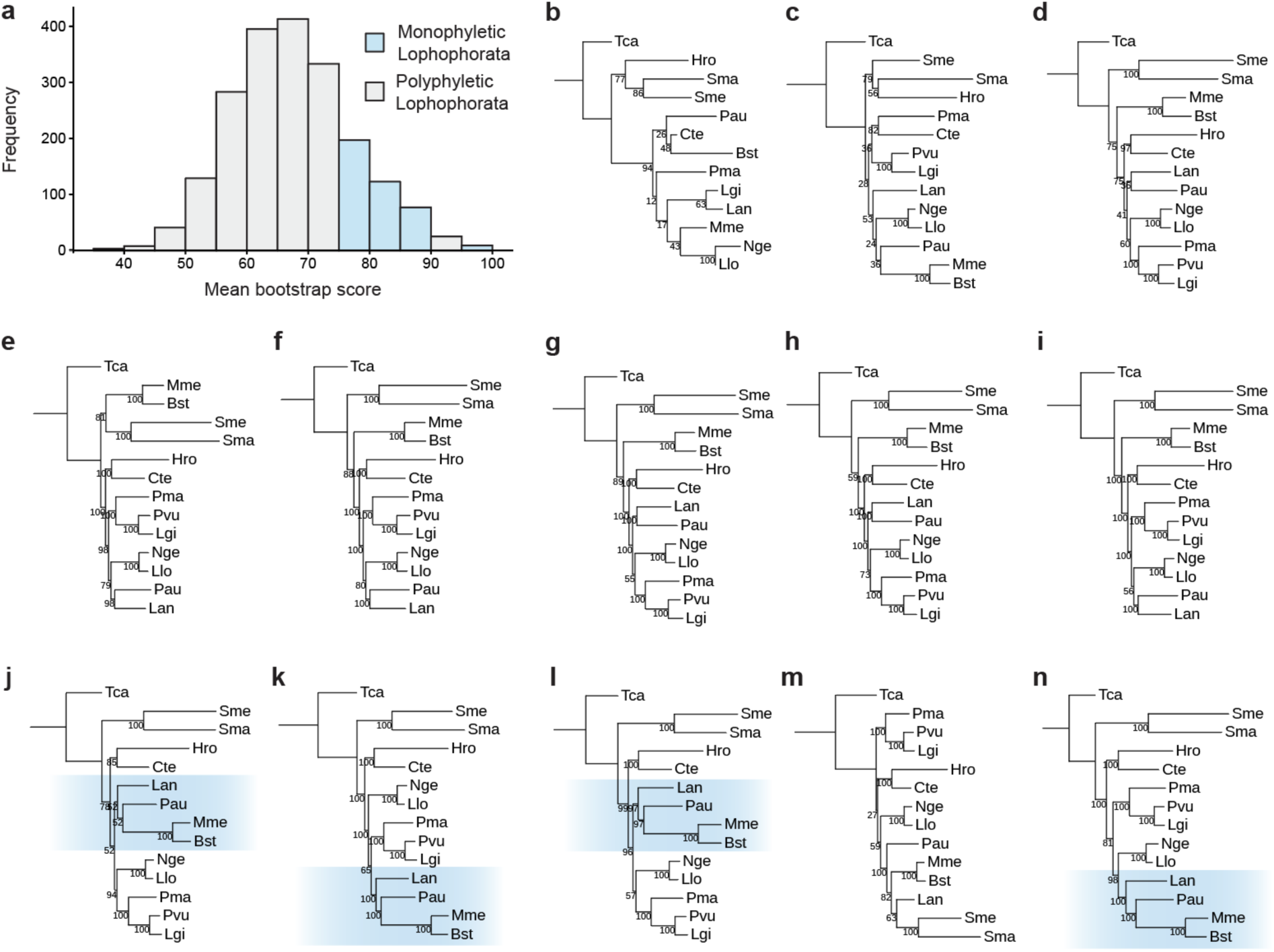
Identification of orthologues with strong phylogenetic signals. **a**, Distribution of mean bootstrap score for species trees built with each of 2,036 orthologues; bin width 5%. Bins are coloured grey or pale blue based on whether a tree made from those orthologues includes a polyphyletic or monophyletic Lophophorata, respectively. **b–n**, Maximum likelihood trees made from orthologues in 5% bins associated with the above analysis. Blue boxes mark the presence of the clade Lophophorata. Trees made from orthologues with higher bootstrap scores tend to support a monophyletic Lophophorata. Trees are made from the following mean bootstrap score bins: **b**, 35–40%; **c**, 40–45%; **d**, 45–50%; **e**, 50–55%; **f**, 55 –60%; **g**, 60–65%; **h**, 65–70%; **i**, 70–75%; **j**, 75–80%; **k**, 80–85%; **l**, 85–90%; **m**, 90–95%; **n**, 95–100%.

**Fig. S5.**
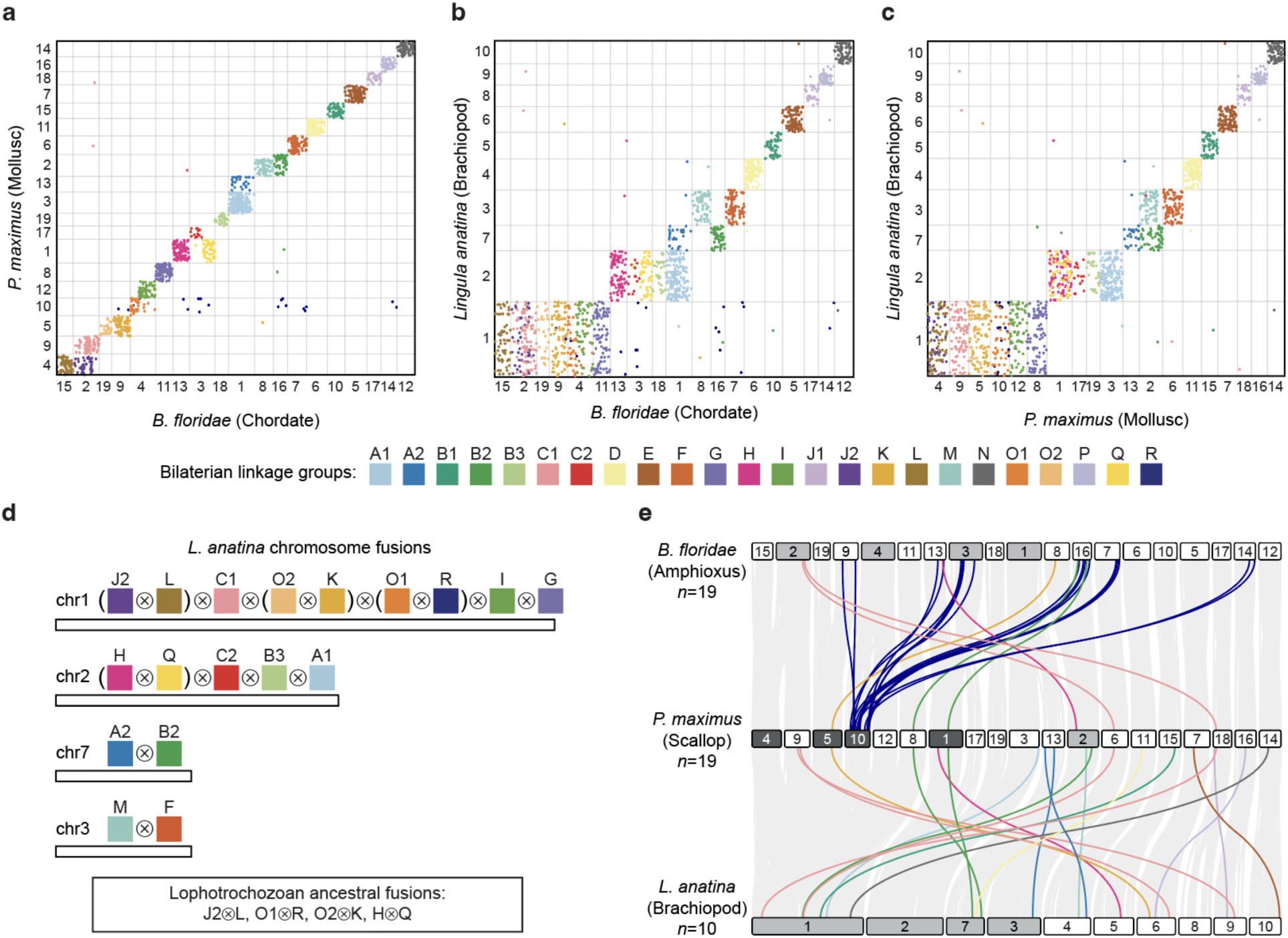
Chromosome-scale gene linkage between *L. anatina*, *P. maximus* and *B. floridae*. Oxford dot plot revealing chromosome-scale gene linkage between the mollusc *P. maximus* and the chordate *B. floridae*. Each axis represents the entire length of the genome of one species. Grey bars separate chromosomes. Each point represents a pair of orthologues, placed by their ordinal position in each genome. Macrosynteny is highly conserved between these two groups, with few chromosome rearrangements. Linkage group R (dark blue) is lost in chordates, and is dispersed around the genome. **b,** Oxford dot plot revealing chromosome-scale gene linkage between the chordate *B. floridae* and the brachiopod *L. anatina*. *L. anatina* chromosome 1 corresponds to six *B. floridae* chromosomes (chr2, 4, 9, 11, 15, 19). *L. anatina* chromosome 2 corresponds to four *B. floridae* chromosomes (chr1, 3, 13, 18). *L. anatina*. chromosome 7 corresponds to two *B. floridae* chromosomes (chr1, 16). *L. anatina*. chromosome 3 corresponds to two *B. floridae* chromosomes (chr7, 8). **c,** Oxford dot plot revealing chromosome-scale gene linkage between the mollusc *P. maximus* and the brachiopod *L. anatina*. *L. anatina* chromosome 1 corresponds to six *P. maximus* chromosomes (chr4, 5, 8, 9, 10, 12). *L. anatina* chromosome 2 corresponds to four *P. maximus* chromosomes (chr1, 3, 17, 19). *L. anatina*. chromosome 7 corresponds to two *P. maximus* chromosomes (chr2, 13). *L. anatina* chromosome 3 corresponds to two *P. maximus* chromosomes (chr2, 6). **d,** Summary of chromosome fusion events inferred from the *L. anatina* genome. The symbol ⊗ represents a fusion-with-mixing event. Fusions that are present not only in brachiopods but also in other lophotrochozoans (annelids, molluscs and nemerteans) are highlighted with brackets. **e,** Macrosynteny between *B. floridae* (Chordata), *P. maximus* (Mollusca), and *L. anatina* (Brachiopoda). To create informative macrosynteny plots, cases where five or fewer genes are translocated to a different ALG are removed from the figures. This plot shows in colour the genes removed from main text Fig. 1b, while pale grey lines represent genes with a conserved ALG, showing relationships between chromosomes. While most chromosomes have few translocations, *P. maximus* chromosome 10 hosts many genes in ALG R that appear to have been translocated in the *B. floridae* genome. This is because, as previously reported, ALG R has been completely lost in *B. floridae*, so remaining genes are dispersed around the genome.

**Fig. S6.**
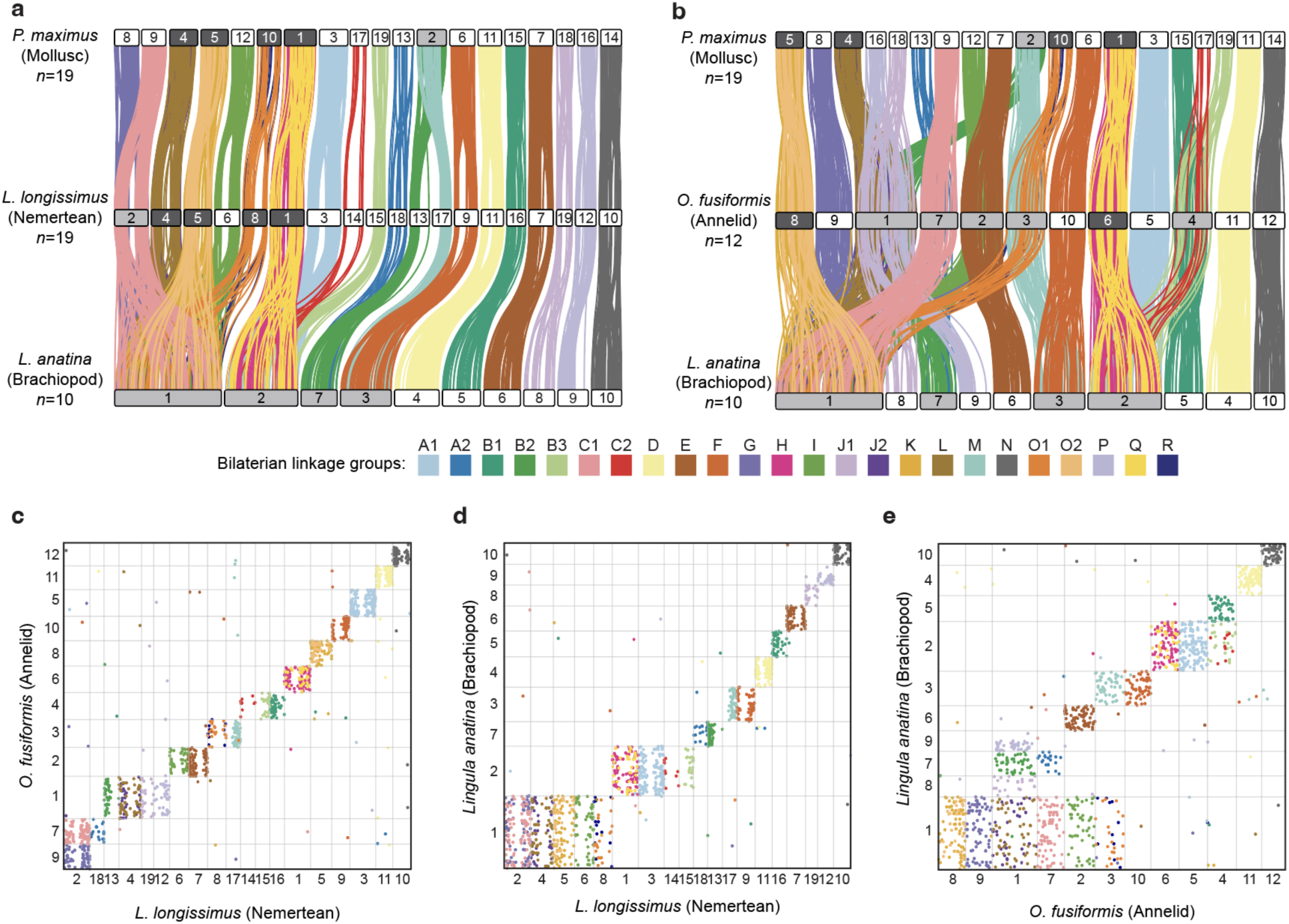
Chromosome-scale gene linkage between *L. anatina*, *L. longissimus*, *O. fusiformis* and *P. maximus*. **a**, Macrosynteny between *P. maximus* (Mollusca), *L. longissimus* (Nemertea) and *L. anatina* (Brachiopoda). Horizontal bars represent chromosomes. Dark grey chromosomes are the products of ancestral lophotrochozoan fusion events. Pale grey chromosomes are the products of lineage-specific fusion events. Vertical lines between chromosomes link orthologous genes and are coloured by bilaterian ALG. *L. longissimus* has undergone minimal genome rearrangements. In addition to the ancestral lophotrochozoan fusion events, it has one additional fusion-with-mixing event, bringing ALGs C1 and G together to form chromosome 2. ALGs C1 and G also occur together on *L. anatina* chromosome 1, but further analysis is required to determine whether these are independent fusions or are the product of a single fusion event in a common ancestor. A striking feature of this plot is the large gene deserts at the centre of the *L. longissimus* chromosomes, which are not observed in any other species. This may be indicative of large centromeric regions in *L. longissimus*. **b**, Macrosynteny between *P. maximus* (Mollusca), *O. fusiformis* (Annelida) and *L. anatina* (Brachiopoda). The *O. fusiformis* genome has undergone several fusion events independent to those occurring in *L. anatina*. In addition to the ancestral spiralian fusion events, chromosomes 1, 2, 3, 4 and 7 are the products of additional fusions. It is notable that only two ALGs (D and N; occupying chromosomes 4 and 10, respectively, in *L. anatina*) have not been involved in a fusion or fission event in the five species in our analysis. We questioned whether these chromosomes harbour specific genes that make fusions evolutionarily deleterious, but gene ontology analysis revealed no significant enrichment of specific gene types on these chromosomes. **c**, Oxford dot plot revealing chromosome-scale gene linkage between the annelid *O. fusiformis* and the nemertean *L. longissimus*. Each axis represents the entire length of the genome of one species. Grey bars separate chromosomes. Each point represents a pair of orthologues, placed by their ordinal position in each genome. **d**, Oxford dot plot revealing chromosome-scale gene linkage between the brachiopod *L. anatina* and the nemertean *L. longissimus*. **e**, Oxford dot plot revealing chromosome-scale gene linkage between the brachiopod *L. anatina* and the annelid *O. fusiformis*.

**Fig. S7.**
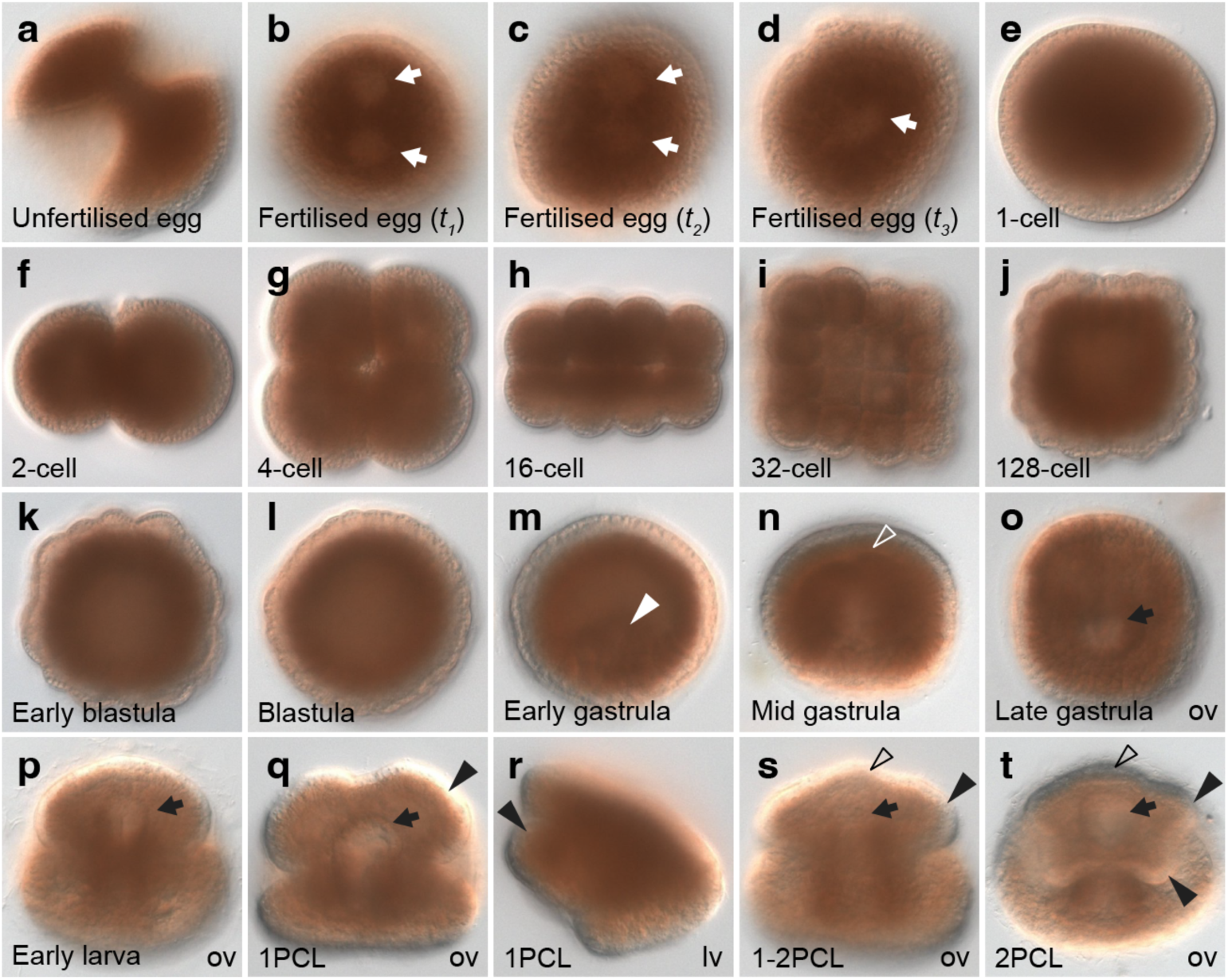
Early embryonic development of the brachiopod *L. anatina*. **a**, Unfertilised eggs typically exhibit an irregular shape. Within 15 min post-insemination, these eggs transform dramatically into a more rounded form. (**b**–**d**) This time sequence, from *t_1_* to *t_3_*, illustrates the pronuclear fusion event post-fertilization. White arrows indicate the pronuclei. (**e**–**t**) Early developmental stages from a single cell to the larval stage. **e**, 1-cell. **f**, 2-cell, **g**, 4-cell, **h**, 16-cell. **i**, 32-cell. **j**, 128-cell. **k**, Early blastula, distinguished by the presence of blastomeres. **l**, Blastula, where a blastoderm is formed. **m**, Early gastrula, starting with invagination (indicated by arrowhead). **n**, The mid gastrula, characterised by the archenteron contacting the ectoderm (shown by a blank arrowhead). **o**, Late gastrula, with the blastopore marked by an arrow. **p**, Early larva, identifiable by the absence of cirri. **q** and **r**, Larval stages with one pair of cirri (1PCL). **s**, Transitional stage from one to two pairs of cirri (1-2PCL), showing tentacle development. **t**, Larval stage with two pairs of cirri (2PCL). Arrows in these stages (**o**–**t**) denote the blastopore, which becomes the mouth area, while the arrowheads and blank arrowheads mark the cirri and tentacles, respectively. ov and lv represent oral and lateral views.

**Fig. S8.**
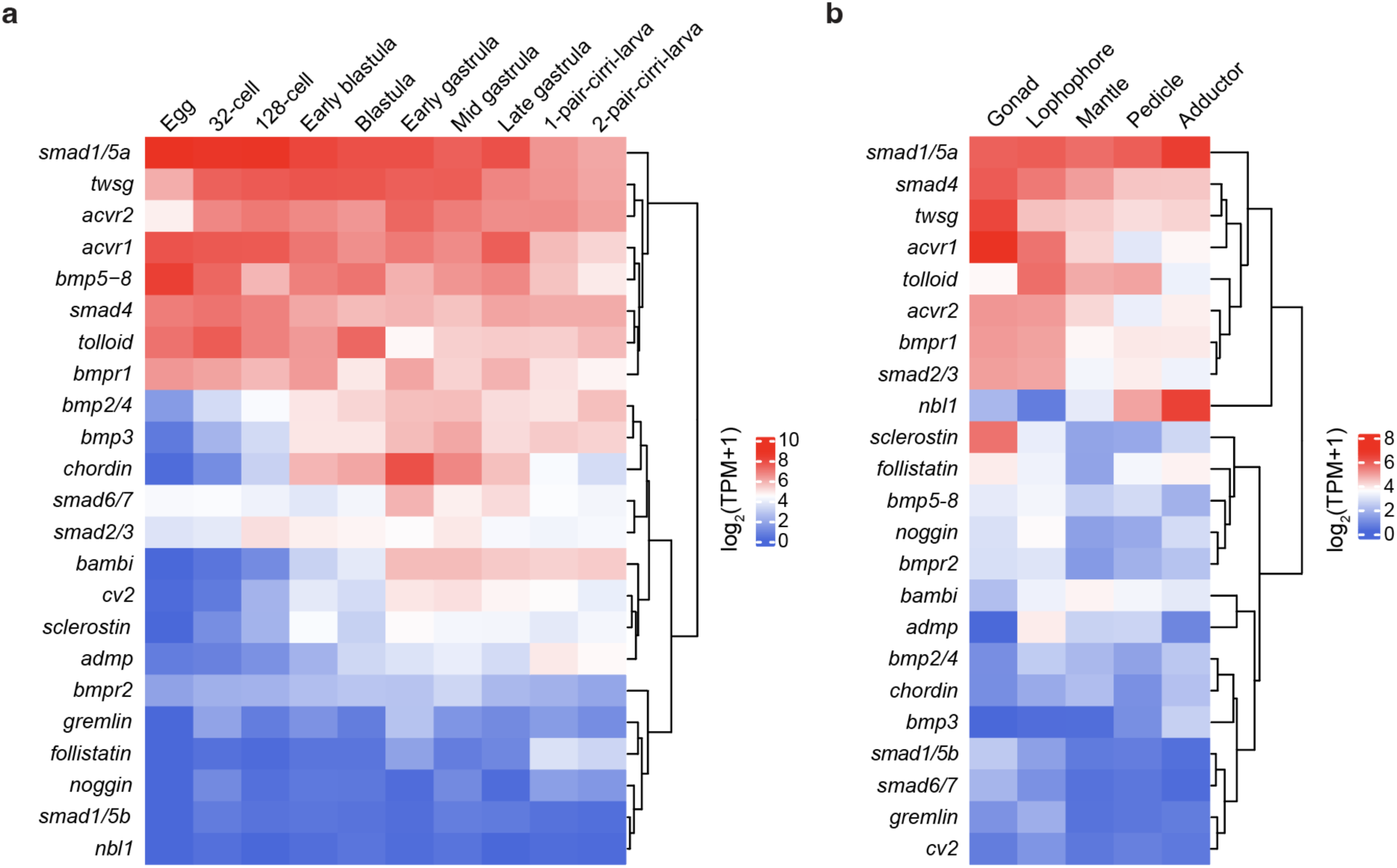
Expression of BMP pathway genes in *L. anatina*. **a**, Expression profiles of BMP signalling ligands, mediators, and modulators during *L. anatina* development. **b**, Expression levels of BMP signalling ligands, mediators, and modulators in five adult *L. anatina* tissues. TPM, transcripts per million.

**Fig. S9.**
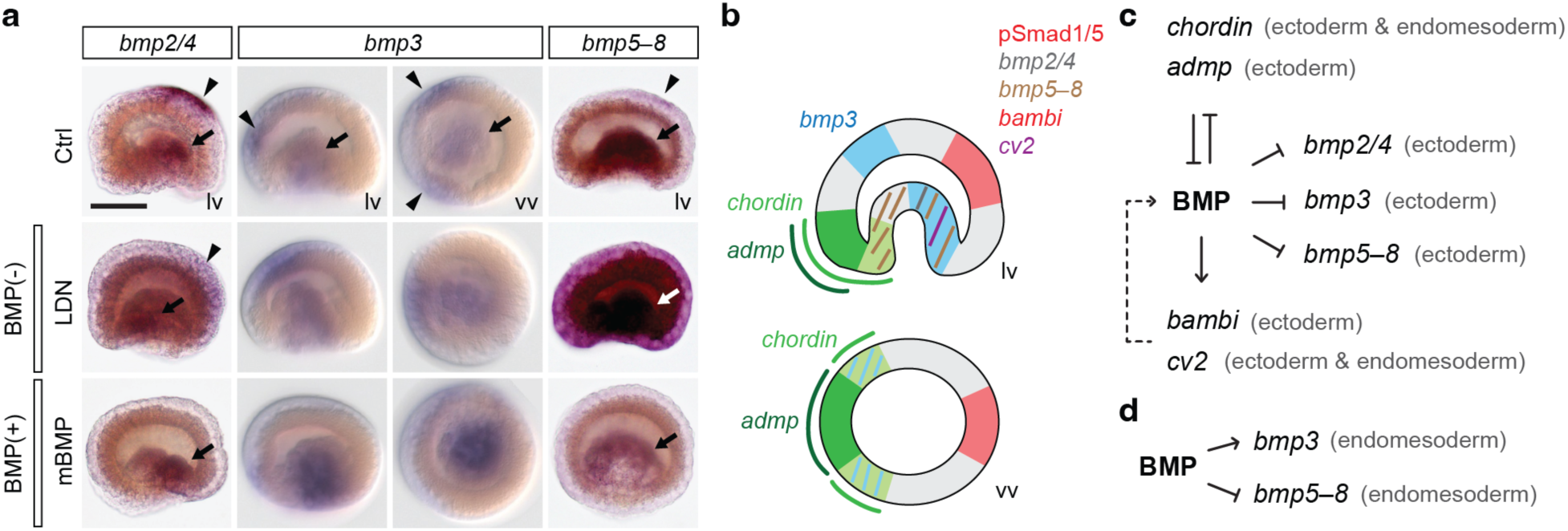
Spatial expression of BMP ligands and regulation of BMP signalling. **a**, Expression profile of BMP ligands in the *L. anatina* gastrula under BMP signalling manipulation. **b**, Cartoon illustration of the expression domains of BMP signalling genes and BMP signals at the late gastrula stage. **c**, Proposed molecular network of BMP signalling genes in the BMP–Chordin axis, where BMP signals may inhibit the expression of BMP ligands in the ectoderm, causing autoinhibition. **d**, Proposed molecular network of BMP signalling genes in the endomesoderm.

