## Supplementary Information for "Brachiopod genome unveils the evolution of the BMP–Chordin network in bilaterian body patterning"

This PDF file includes:

Supplementary Figs. 1 to 6
Legends for Supplementary Tables 1 to 37

Other supporting materials for this manuscript include:

Supplementary Tables 1 to 37

### Supplementary Figs. 1 to 6

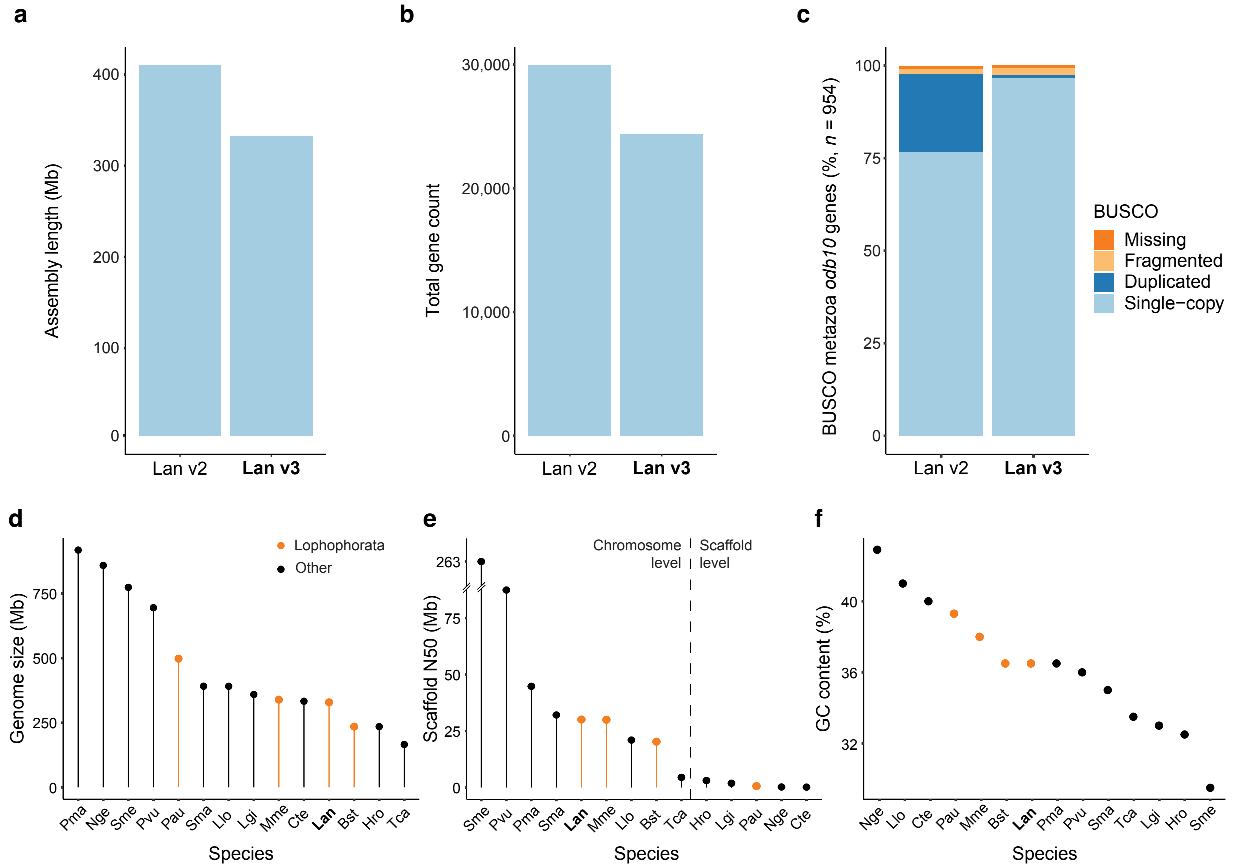


**Supplementary Fig. 1 | Comparison between chromosome-level (this study) and draft assembly of the *L. anatina* genome. a**, Total assembly lengths for the current (Lan v3) and previous (Lan v2) *L. anatina* assemblies (329 Mb versus 406 Mb). **b**, Total gene counts for current and previous *L. anatina* assemblies (24,330 versus 29,907). **c**, BUSCO Metazoa *odb10* for the current and previous *L. anatina* assemblies (0.9% duplication versus 20.8% duplication). Lower duplication, total gene count and shorter assembly lengths of the current assembly versus previous suggest false heterotype duplications were present in the previous assembly but are corrected in the current one. **d**, Genome size of selected spiralian assemblies. Species within the clade Lophophorata are highlighted in orange. **e**, Scaffold N50 of selected spiralian assemblies. Species within the clade Lophophorata are highlighted in orange. **f**, GC content of selected spiralian assemblies. Species within the clade Lophophorata are highlighted in orange. Abbreviations: Bst, *Bugulina stolonifera*; Cte, *Capitella teleta*; Hro, *Helobdella robusta*; Lan, *Lingula anatina*; Lgi, *Lottia gigantea*; Llo, *Lineus longissimus*; Mme, *Membranipora membranacea*; Nge, *Notospermus geniculatus*; Pau, *Phoronis australis*; Pma, *Pecten maximus*; Pvu, *Patella vulgata*; Sma, *Schistosoma mansoni*; Sme, *Schmidtea mediterranea*; Tca, *Tribolium castaneum*.

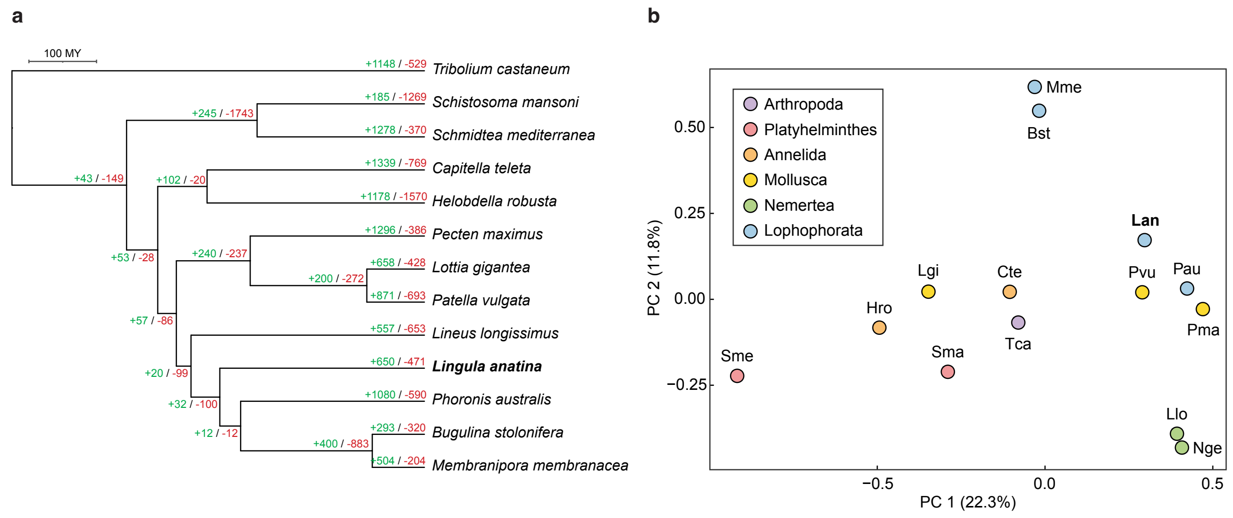


**Supplementary Fig. 2 | Gene family evolution in spiralians. a**, Output of CAFE analysis. Figures in red and green are estimated gene gains and losses, respectively. The brachiopod *L. anatina* has a relatively conservatively evolving gene content compared to other Lophophorata members: phoronids (*Phoronis australis*) have a large number of gene gains, while bryozoans (*Bugulina stolonifera* and *Membranipora membranacea*) have a high rate of gene loss. **b**, Principal component analysis (PCA) of spiralian gene content. OrthoFinder was used to create an orthogroup count matrix. The gene content of *L. anatina* (Lan) is most similar to that of phoronids (Pau) and molluscs (Pvu and Pma). The platyhelminths *Schmidtea mediterranea* and *Schistosoma mansoni* (Sme and Sma), the annelid *Helobdella robusta* (Hro) and the bryozoans *B. stolonifera* and *M. membranacea* all have extensive gene loss. The separate grouping of the two bryozoans away from the platyhelminths and annelid suggest different gene sets have been lost in these two groups. Abbreviations as in Supplementary Fig. 1.

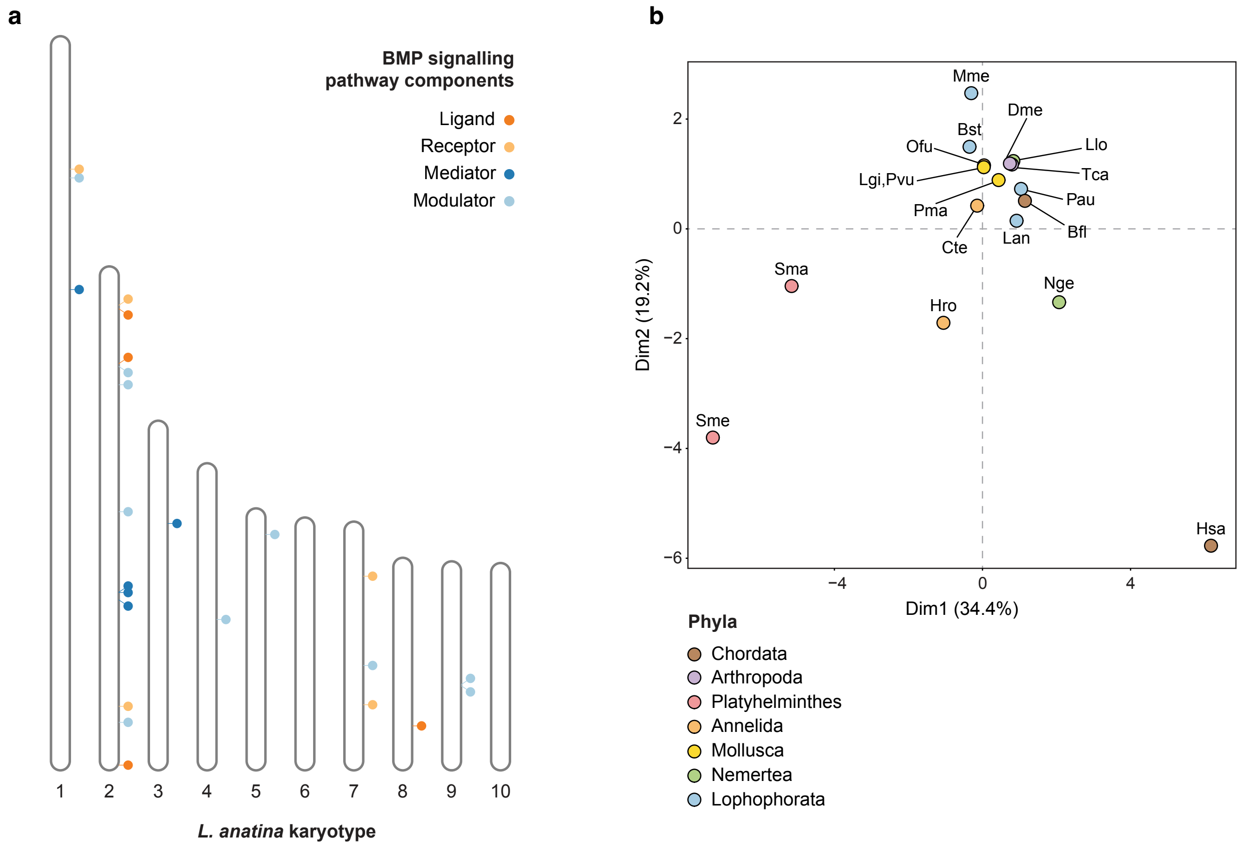


**Supplementary Fig. 3 | Evolution of BMP signalling pathway components. a**, Locations of BMP signalling pathway-related genes (circles) in the *L. anatina* genome. Vertical bars represent chromosomes. **b**, Principal component analysis (PCA) using species' BMP gene repertoires. Abbreviations as in Supplementary Fig. 1.

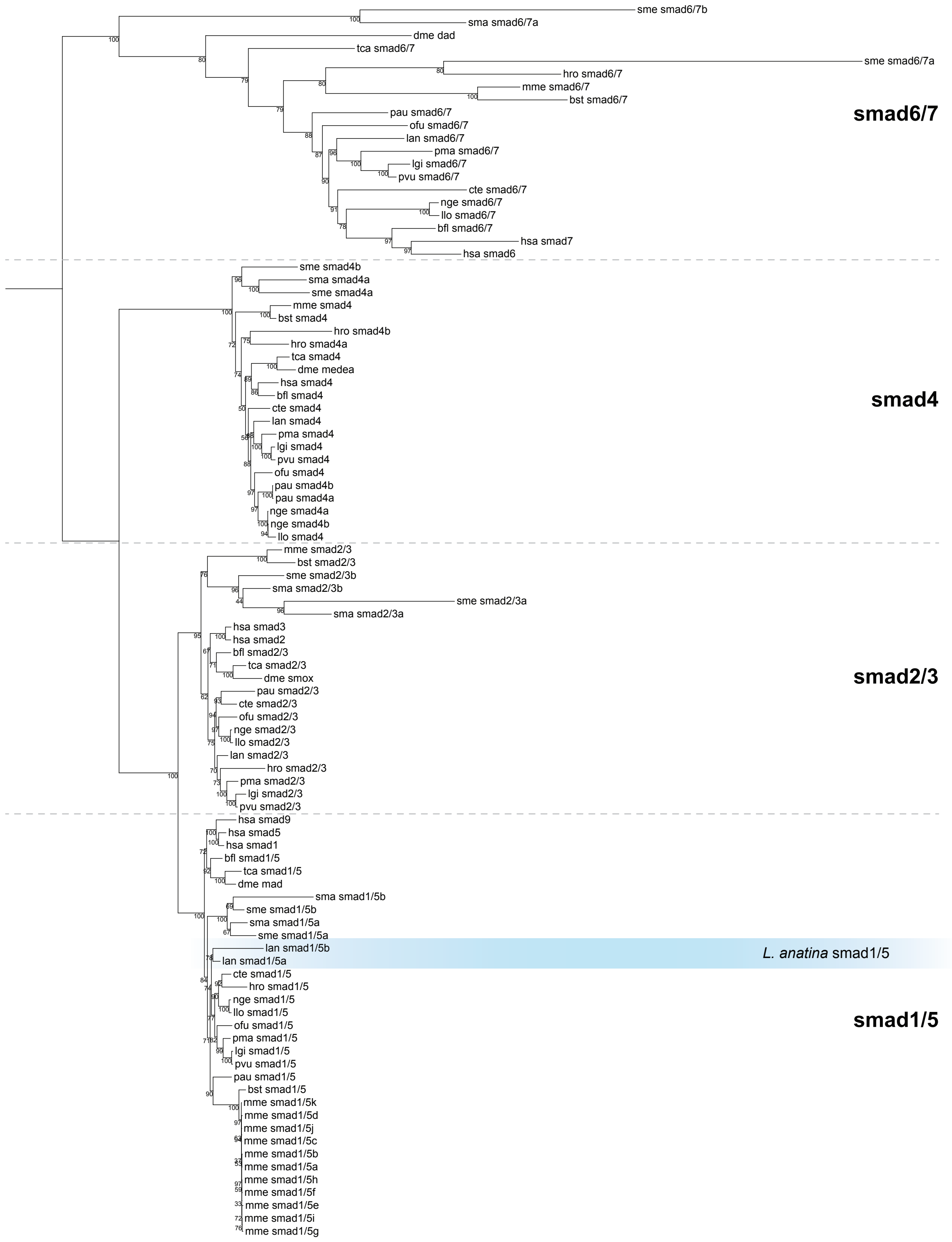


**Supplementary Fig. 4 | Tree of Smad proteins identified in this work.** Tree was produced using the maximum likelihood method (model LG+F+R6) with 1000 bootstrap replicates. *L. anatina* (lan) has a lineage-specific smad1/5 duplication. The bryozoan *M. membranacea* has 11 copies of *smad1/5*. Abbreviations as in Supplementary Fig. 1.

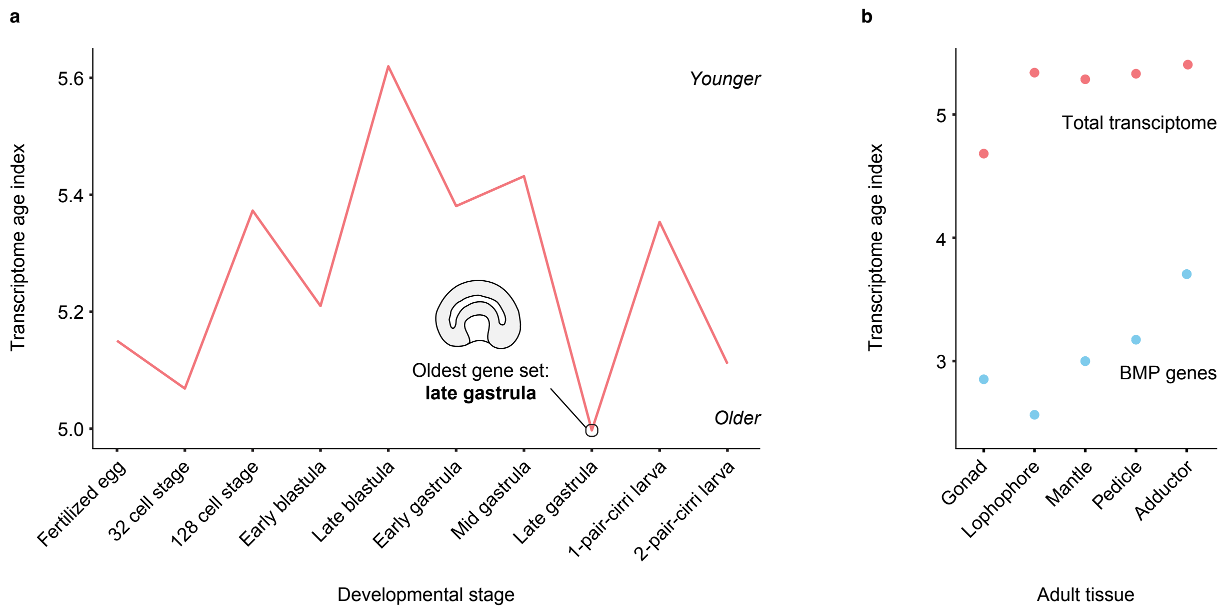


**Supplementary Fig. 5 | Transcriptome age index (TAI) in *L. anatina* tissues. a**, TAI of ten stages of *L. anatina* development, from the fertilised egg to the larval stage. The youngest gene set occurs at the late blastula, while the oldest is observed at the late gastrula. The late gastrula stage is a key time for axis specification, and has the oldest transcriptome. **b**, TAI of six adult tissues. For the total transcriptome, the oldest gene set is observed in ovary tissue and youngest in adductor muscle. The TAI of BMP-related genes is considerably lower than that of the overall transcriptome, reflecting their ancient evolutionary origins.

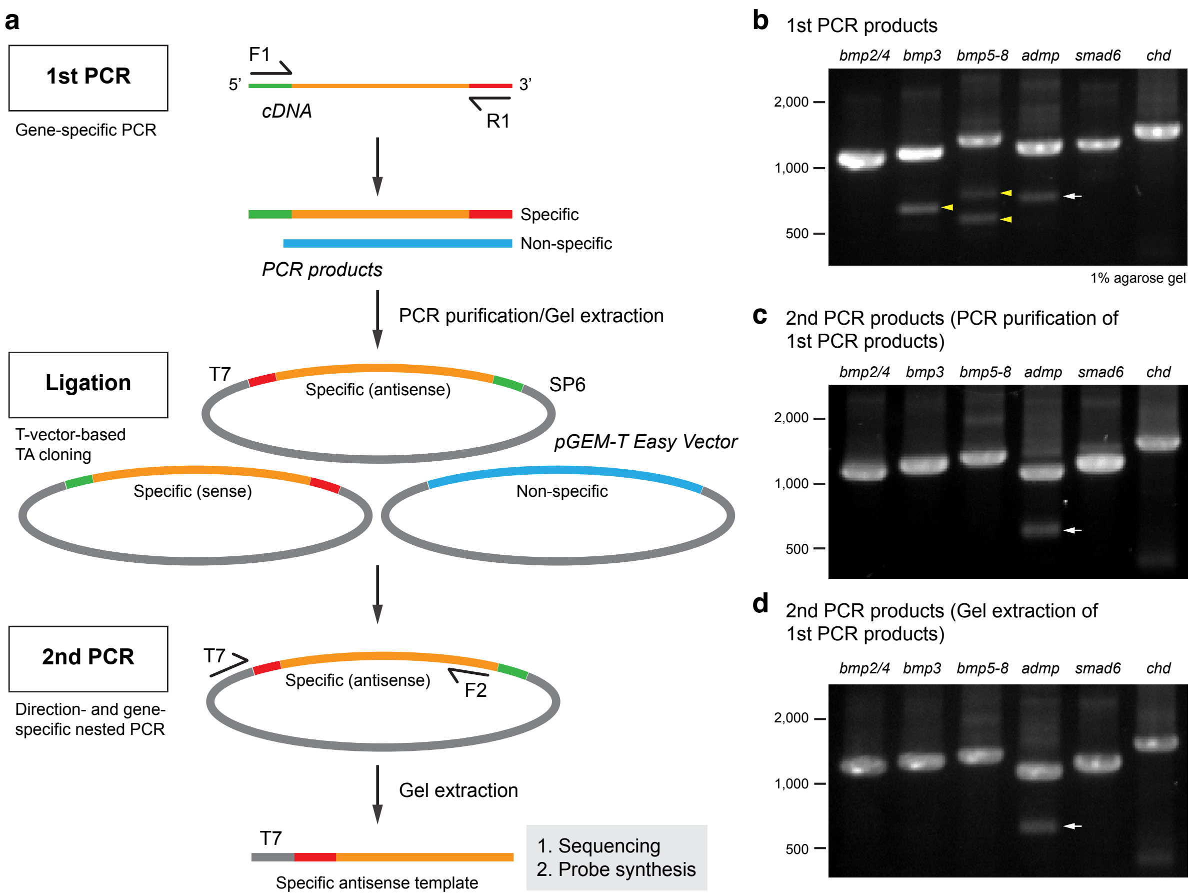


**Supplementary Fig. 6 | Bacterial-cloning-free synthesis of RNA probes. a**, Schematic of the process to create specific antisense DNA templates using T-vector-based ligation and nested PCR. The main products from the second PCR, which are of the predicted size, are extracted from a gel for sequencing and RNA probe synthesis. **b**, Agarose gel electrophoresis results of the first PCR products, with examples of BMP signalling components. Yellow arrowheads highlight non-specific bands. A white arrow points to a minor band of unknown origin (potentially arising from DNA secondary structures or non-specific internal insert regions). **c**, Agarose gel electrophoresis of the second PCR products, created using the first PCR products after PCR purification in the ligation reactions. **d**, Agarose gel electrophoresis of the second PCR products, obtained using ligation reactions from the first PCR products followed by gel extraction. Note that non-specific bands are eliminated after the nested (second) PCR, and no significant differences are observed between PCR purification and gel extraction when using the first PCR products for ligation.

**Legends for Supplementary Tables 1 to 37**

**Supplementary Table 1 |** Sequencing statistics for the *Lingula anatina* genome.

**Supplementary Table 2 |** Hi-C-assisted genome scaffolding.

**Supplementary Table 3 |** Individual scaffold statistics for the *L. anatina* genome.

**Supplementary Table 4 |** Annotation of repeats in the *L. anatina* genome.

**Supplementary Table 5 |** Assembly statistics for the *L. anatina* genome.

**Supplementary Table 6 |** *L. anatina* gene annotation.

**Supplementary Table 7 |** Protein annotation with InterProScan.

**Supplementary Table 8 |** Protein annotation with KEGG orthology implemented in KofamScan.

**Supplementary Table 9 |** Protein annotation with eggNOG.

**Supplementary Table 10 |** Orthologues of *L. anatina* proteins in a mollusc (*P. vulgata*, common limpet) and a chordate (*Homo sapiens*, human) identified with OrthoFinder.

**Supplementary Table 11 |** Input dataset for CAFE 5 gene family evolution analysis. Abbreviations as in Supplementary Fig. 1.

**Supplementary Table 12 |** Genomes used for phylogenetic and comparative genomic analyses.

**Supplementary Table 13 |** Lophotrochozoan BMP gene repertoires. Abbreviations as in Supplementary Fig. 1.

**Supplementary Table 14 |** *L. anatina* BMP pathway protein sequences.

**Supplementary Table 15 |** Chromosome ancestral linkage group (ALG) assignments for macrosynteny analysis.

**Supplementary Table 16 |** Chromosome rearrangements in study species.

**Supplementary Table 17 |** Conserved associations of developmental genes with ALGs.

**Supplementary Table 18 |** Chi-square test for conserved ALG associations of BMP genes.

**Supplementary Table 19 |** Chi-square test for conserved ALG associations of Wnt genes.

**Supplementary Table 20 |** Expression of BMP pathway genes (TPM) during *L. anatina* embryonic development.

**Supplementary Table 21 |** Summary of BMP signalling manipulation experiments. Visualisation presented as main text Fig. 3a.

**Supplementary Table 22 |** Summary of RNA-seq samples from BMP signalling manipulation experiments.

**Supplementary Table 23 |** Correspondence of genome-based gene models to transcriptome.

**Supplementary Table 24 |** Gene expression (TPM) in BMP signalling manipulation experiments.

**Supplementary Table 25 |** Gene ontology analysis for genes upregulated by BMP signalling at the late gastrula stage.

**Supplementary Table 26 |** Gene ontology analysis for genes downregulated by BMP signalling at the late gastrula stage.

**Supplementary Table 27 |** Gene ontology analysis for genes upregulated by BMP signalling at the larval stage.

**Supplementary Table 28 |** Gene ontology analysis for genes downregulated by BMP signalling at the larval stage.

**Supplementary Table 29 |** Transcriptome age index analysis of *L. anatina* developmental stages and adult tissues.

**Supplementary Table 30 |** Transcriptome age index analysis of *L. anatina* late gastrula embryos and larvae under conditions of BMP signalling manipulation.

**Supplementary Table 31 |** *t*-tests for differences in transcriptome age index score between BMP signalling manipulation conditions.

**Supplementary Table 32 |** Gene expression in *L. anatina* adult tissues.

**Supplementary Table 33 |** Gene expression in *L. anatina* developmental stages.

**Supplementary Table 34 |** Published RNA-seq datasets used to annotate the *L. anatina* genome with BRAKER.

**Supplementary Table 35 |** Ancestral linkage group associations of genes across bilaterians.

**Supplementary Table 36 |** Gene ages (‘phylostrata’) in the *L. anatina* genome estimated with GenEra.

**Supplementary Table 37 |** Supplementary genomes added to GenEra.
